# Capturing the emergent dynamical structure in biophysical neural models

**DOI:** 10.1101/2024.10.21.619355

**Authors:** Borjan Milinkovic, Lionel Barnett, Olivia Carter, Anil K. Seth, Thomas Andrillon

## Abstract

Complex neural systems can display structured emergent dynamics. Capturing this structure remains a significant scientific challenge. Using information theory, we apply *Dynamical Independence* (DI) to uncover the emergent dynamical structure in a minimal 5-node biophysical neural model, shaped by the interplay of two key aspects of brain organisation: integration and segregation. In our study, functional integration within the biophysical neural model is modulated by a global coupling parameter, while functional segregation is influenced by adding dynamical noise, which counteracts global coupling. DI defines a dimensionally-reduced *macroscopic variable* (e.g., a coarse-graining) as emergent to the extent that it behaves as an independent dynamical process, distinct from the micro-level dynamics. We measure dynamical dependence (a departure from dynamical independence) for macroscopic variables across spatial scales. Our results indicate that the degree of emergence of macroscopic variables is relatively minimised at balanced points of integration and segregation and maximised at the extremes. Additionally, our method identifies to which degree the macroscopic dynamics are localised across microlevel nodes, thereby elucidating the emergent dynamical structure through the relationship between microscopic and macroscopic processes. We find that deviation from a balanced point between integration and segregation results in a less localised, more distributed emergent dynamical structure as identified by DI. This finding suggests that a balance of functional integration and segregation is associated with lower levels of emergence (higher dynamical dependence), which may be crucial for sustaining coherent, localised emergent macroscopic dynamical structures. This work also provides a complete computational implementation for the identification of emergent neural dynamics that could be applied both in silico and in vivo.

**Author summary:** Understanding how complex neural systems give rise to emergent macroscopic patterns is a central challenge in neuroscience. Emergence, where macroscopic structures appear from underlying microscopic interactions, plays a crucial role in brain function, yet identifying the specific dynamics involved remains elusive. Traditionally, methods have quantified the extent of emergence but have struggled to pinpoint the emergent dynamical structure itself. In this study, we develop and apply a method, based on a quantity called Dynamical Independence (DI), which simultaneously captures the extent of emergence and reveals the underlying dynamical structure in neurophysiological data. Using a minimal 5-node biophysical neural model, we explore how a balance between functional integration and segregation—two key organisational principles in the brain—affects emergent macroscopic dynamics. Our results show that a finely balanced system produces highly localised, coherent macroscopic structures, while extreme deviations lead to more distributed, less localised dynamics. This work provides a computational framework for identifying emergent dynamical structure in both theoretical models and potentially in empirical brain data, advancing our understanding of the brain’s complex organisation across higher-order scales.

## Introduction

The co-existence of functional integration and segregation between distinct brain regions has been argued to support perceptual and cognitive states [1–6]. Typically, for a system to be fully integrated, strong interactions between microlevel constituents must be reinforced and drive the dynamics; tending towards complete order through global coupling [7–9]. Conversely, for a system to be fully segregated, local fluctuations must dominate global influences induced by coupling, fracturing functional connectivity and segregating the microlevel constituents [10, 11]. Dynamical noise has previously been employed as a way to functionally segregate variables with known anatomical connectivity [10, 12, 13], and is used in dynamical systems theory as a way of reducing the effect of the coupling parameter in state-variables [9, 14]. Yet, at some balanced point(s) between complete functional integration and segregation, dynamics emerge as highly organised patterns of activity [15, 16]. A pressing issue in computational neuroscience remains the quantification and identification of these emergent dynamics.

Neural complexity measures, some of which peak at a balance between functional integration and segregation, have been successfully used as biomarkers of conscious states [17–23]. More generally, the co-existence of these two aspects of brain organisation has been associated with *criticality* in complex systems [6, 24]. Related to criticality [25], empirical measures of *integrated information* also peak at a balanced point between functional integration and segregation in brain data [18, 26–29]. This previous work highlights an important and unresolved challenge: to precisely characterise how the interplay between functional integration and segregation mediates the emergent macroscopic *dynamical structure* of the underlying brain network. Simply, to not only quantify when emergent dynamics occur, but also to be able to describe them. We address this challenge by using a minimal 5-node biophysical neural model and leveraging an information-theoretic measure—called *Dynamical Independence* (DI)— of emergence [30]. Specifically, we set out to (i) characterise how the modulation of functional integration and segregation impacts the emergent dynamics and (ii) identify the underlying macroscopic dynamical structure across spatial scales. We thus provide a complete computational implementation to capture emergent dynamical structures, which could be applied to both in vivo and in silico recordings.

In this paper, we examine the *emergent dynamical structure* by focusing on how macroscopic dynamics are distributed across specific microlevel nodes. Specifically, we quantify the extent to which each microlevel node contributes to the overall macroscopic process, thereby highlighting the *localisation* of these dynamics.

In general, capturing the organisation of brain activity at a macroscopic level has been approached through various methods, including absolute signal correlations [31], clustering analysis [32], small-worldness and graph theory [33], phase synchronisation [34], metastability [35, 36], and more recently, eigenmode decomposition techniques [37–39]. However, while these methods often capture functional structures, they do not explicitly reveal *emergent* macroscopic dynamical structure arising from microlevel interactions. Given that the brain’s activity fluctuates between a variable degree of the co-existence of integration and segregation, the information flow between regions may transition from predictable and structured (integrated) to unpredictable and stochastic (segregated) [40]. Information theory [41], which quantifies how one node in the network predicts the behaviour of another [42–45], emerges as a natural candidate for uncovering macroscopic dynamical structure from microlevel interactions [30, 46, 47].

Recent information-theoretic approaches to emergence show considerable promise in revealing the capacity for higher-order interactions [48–51]. Though these approaches align with capturing the capacity for emergence in brain networks, they do not explicitly consider the relation between functional integration and segregation and its effect on the emergent dynamical structure. Further, with the exception of [51], when applied to larger systems, measures based on decomposing information are approximated from pairwise interactions between microlevel variables alone, and do not capture macroscopic organisation across the full array of possible spatial scales.

To understand the relation between emergent dynamical structure and functional integration and segregation in neural systems a metric is needed that simultaneously captures i) coarse-grained macroscopic patterns that represent the *dynamical structure* of the system’s interactions and ii) measures the degree to which the coarse-grainings are to be considered as *emergent* variables with respect to the microlevel constituents.

The problem can be framed as an optimisation task where macroscopic (coarse-grained) variables across spatial scales are identified to best describe the underlying stochastic dynamical process. This approach highlights how brain activity can manifest organisation at the macroscopic level, revealing patterns not evident from the microlevel perspective alone. This data-driven approach would be a significant advance over existing complexity-based approaches in neuroscience, offering deeper insights into how emergent structure arises in neural processes—from the ground, up.

Consequently, we apply DI to specifically identify emergent macroscopic variables in complex dynamical systems across all spatiotemporal scales. DI captures macroscopic coarse-grainings of the system’s dynamics whereby the self-prediction of the future state of the coarse-grained macroscopic variable is not significantly improved by knowledge of the historical past of the microlevel dynamics. DI therefore identifies macroscopic coarse-grainings of the dynamics *generated by* the interactions between the microlevel constituents but that appear to be independent of them. Because DI can be applied directly to continuous-valued random variables, it is ideally suited for revealing the emergent dynamical structure in neurophysiological data.

To apply Dynamical Independence (DI) to biophysical neural models, we start by simulating a brain network model with five nodes. Each node’s local dynamics are defined by the Stefanescu-Jirsa 3D (SJ3D) neural mass model, which simulates the neural population activity of brain regions capable of spike-burst behavior [52, 53]. This model was specifically chosen given recent studies proposing spike-bursts as a neural mechanism for conscious processing [54, 55].

While this model is not a large-scale whole-brain model informed by MRI/DTI data, it is nevertheless a complex biophysical model that incorporates biologically realistic local dynamics, global coupling, dynamical noise, and temporal delays. The term *biophysical neural model* is preferred to emphasise that, while it is not a full representation of whole-brain dynamics, it remains grounded in biological realism by displaying local activity attributed to neural populations [53], and system-system wide activity governed by three parameters known to influence whole-brain activity; global coupling, dynamical noise, and temporal delay [56–58]. This study uses the model as a means to validate the approach in a controlled, albeit simplified, setting, acknowledging that it does not represent the full complexity of whole-brain dynamics but is still informed by the principles that govern such systems.

In the brain network model, functional integration is modulated by the global coupling parameter, influencing the strength of connections between regions and affecting local dynamics [9, 59]. Conversely, functional segregation is influenced by dynamical noise, which reduces the signal-to-noise ratio, thereby limiting the impact of global coupling on local dynamics [9, 12, 56]. Biophysically, this injected noise can be interpreted as representing the influence of unobserved exogenous inputs to the network, which are not explicitly modeled but affect the system’s behaviour.

We then minimised the dynamical dependence (DD) of macroscopic variables across varying degrees of functional integration and segregation. Our results show that when the coexistence of integration and segregation are balanced, the DD of macroscopic variables is higher compared to other parameter regime conditions, indicating that macroscopic dynamics are less emergent and more dependent on the underlying micro-level dynamics. This seems counterintuitive because measures of neural and dynamical complexity typically peak when integration and segregation are balanced [1, 18, 26]. However, at these balanced points, the emergent dynamical structure—defined by the *localisation* of macroscopic variables—is maximised. In our context, localisation refers to how closely the emergent macrovariables align with specific axes of the microscopic state space—each axis corresponding to a neural source or node. A macrovariable is considered localised when it closely aligns with one or a few of these axes, meaning it primarily captures the dynamics of specific microlevel components rather than being a distributed combination of many nodes. This alignment indicates that specific microlevel nodes contribute predominantly to the macroscopic dynamics. Conversely, deviating from these balanced points leads to a loss of localisation, where the contributions of microlevel nodes to the macroscopic variables become more distributed. These findings suggest that when functional integration and segregation are finely balanced, the macroscopic structure exhibits lower degrees of emergence in terms of dynamical dependence but simultaneously achieves maximal organisation into distinct localised contributions from the micro-level nodes.

## Models and methods

### Theory

#### Coarse-graining and macroscopic variables

Consider a discrete-time multivariate stochastic process *X* taking values *X*_*t*_ in a state space 𝒳, which we will refer to as the microlevel or microscopic scale of the system, where *t* ∈ ℤis the discrete time index. (We generally set specific state values in lower case, and random variables in upper case. When referring to a stochastic process as a whole, we write just *X*.) Generally, 𝒳 will have some additional mathematical structure, e.g., topological, metric, differentiable, linear, etc. Later we shall specialise to the case where 𝒳= ℝ^*N*^, i.e., *N* -dimensional Euclidean space with the usual metric and linear vector space structure. In that case, we use standard vector-matrix notation and set vector quantities in bold type; so the microscopic system becomes a multivariate vector process ***X*** defined by 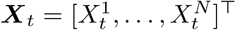 with superscripts indexing components.

Now consider a surjective, structure-preserving mapping *M* : 𝒳 → 𝒴 from the microscopic state-space 𝒳to a *macroscopic* state space 𝒴 Surjectivity means that for every *y* ∈ 𝒴, there exists at least one element *x* ∈ 𝒳 such that *y* = *M* (*x*). We refer to such a mapping as a *coarse-graining*. A coarse-graining effects a partitioning of the microscopic state space as a disjoint union: 𝒳= _*y*∈𝒴_ *M* ^−1^(*y*). However, distinct coarse-grainings *across scales* do not necessarily partition the dimensions of the microscopic state space into non-overlapping sets. Meaning that the dimensions *X*^*n*^ ∈ 𝒳 could belong to multiple coarse-grainings. This also results in the possibility that coarse-grainings can be nested.

The macroscopic state space 𝒴 will typically be of smaller cardinality or dimension—which we refer to as the *scale* of the coarse-graining—so coarse-grainings may be considered as dimensional reductions. The coarse-graining *M* naturally maps the microscopic process *X* to the *macroscopic* process (or macroscopic variable) *Y* defined by *Y*_*t*_ = *M* (*X*_*t*_); this will in general entail a loss of information. In the Euclidean case 𝒳 = ℝ^*N*^, 𝒴 = ℝ^*n*^ with 0 *< n < N*, in order to preserve the vector space structure we consider only *linear* coarse-grainings, so that *M* becomes an *n × N* full-rank^1^ matrix operator, and we write macrovariables as ***Y*** _*t*_ = *M* ***X***_*t*_.

#### The DI framework

DI is a data-driven information-theoretic principle aimed at the identification of emergent coarse-grained macroscopic variables from discrete-valued or continuous-valued time-series data [30]. Intuitively, a macroscopic variable *Y* given by *Y*_*t*_ = *M* (*X*_*t*_) is dynamically independent if—notwithstanding its deterministic dependence on the microscopic process—it behaves like a dynamical process in its own right, following its own dynamical laws, distinct from the laws governing the dynamics of the microlevel variable *X*. Note that in general an arbitrary macroscopic variable will *not* have the property of dynamical independence from the micro-level base.

DI is defined in a *predictive* sense: *Y* is dynamically independent of *X* if knowledge of the history of *X* does not enhance prediction of *Y* beyond the extent to which *Y* already self-predicts. Discovering the macroscopic variables, *Y*, may be framed as an optimisation problem; namely to minimise,across all coarse-grainings, the objective function of *dynamical dependence* (DD), an information-theoretic measure of *departure* from dynamical independence for macroscopic variables:

##### Definition 1.

**Dynamical Dependence** is defined as the *transfer entropy* [60, 61]—a *nonparametric measure of information flow—from the historical past of the microscopic process X* to the present state of the macroscopic process, *Y* at time *t*:^2^

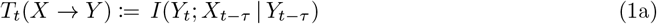

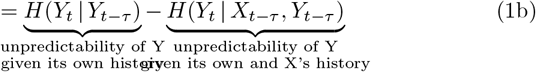

Here, we impose a supervenience relation, which asserts that:

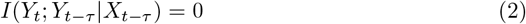

This indicates that there is no additional information in the macroscopic process beyond what is already captured in the history of the microscopic process.

Consequently, within the DI framework, there is no room for additional information on the macroscopic process in the DI framework. Note that a coarse-grained macroscopic variable *Y*_*t*_ = *M* (*X*_*t*_) trivially satisfies (2).

If the process *X* (and hence also *Y*) is *stationary* —i.e., its statistical structure does not change over time—then the DD is not time-dependent, and we drop the subscript *t*. We assume stationarity for all processes from here on. In practice, to compute dynamical dependence from neurophysiological data such as that simulated by biologically-plausible brain network models (see next Section), we employ an approximation method based on linear state-space modelling (see Section S2 Appendix: Linear state-space modelling for details).

Thus a macrovariable is dynamically independent if and only if (iff) its dynamical dependence on the microlevel vanishes identically; i.e., its capacity to predict its future based only on its own history is not enhanced by knowledge of the history of the microscopic process. We define this with a transfer-entropic identity:

##### Definition 2.

A macrovariable *Y* given by the coarse-graining *Y*_*t*_ = *M* (*X*_*t*_) is **dynamically independent** of the microscopic process *X* iff

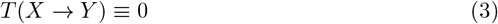

This formalises the intuition of an emergent macrovariable as one which behaves as a dynamical process in its own right, independently of the micro-level dynamics. In the language of Bertschinger *et al*. [62, 63], *the macroscopic variable is informationally (or dynamically) closed* with respect to the microscopic dynamics. Dynamical independence encapsulates the specific notion of emergence discussed in this article, and dynamical dependence stands as a quantitative information-theoretic measure of the degree of (non-)emergence.

In general, given a microscopic process *X*, there may well be *no* perfectly dynamically-independent macroscopic variables (i.e., variables for which Eq 3 holds identically) at some, or indeed at any, spatial scale. Even if there were, in an empirical scenario with finite data, due to the statistical nature of DI, it is in principle impossible to establish that a macroscopic variable at some given scale is in fact perfectly dynamically-independent. Instead, we search for macrovariables with *minimal* DD—i.e., macroscopic variables with a significant degree of emergence. In practice, at any given scale, we attempt to identify macroscopic variables that minimise the DD over the space of all possible structure-preserving coarse-grainings at that scale. Importantly, in an empirical scenario, *minimal* DD may be quantified in a principled manner: namely, that at some predefined significance level, and accounting appropriately for multiple hypotheses, we cannot reject the null hypothesis that an optimised DD value is zero.

For non-trivial problems like the biophysical neural model considered in this study, optimisation is generally analytically intractable, so we are bound to deploy numerical methods. DD minimisation—which will generally take the form of multiple optimising runs with random initial configurations—will, furthermore, be limited by time and computational resources, so will not be sure to guarantee that globally minimal-DD macrovariables are identified. Indeed, it turns out that numerical optimisation runs are in practice likely to terminate at local suboptima of the DD objective function. We therefore adopt the following pragmatic criterion for the discovery of emergent macroscopic variables:

##### Definition 3.

**Emergent** *n***-macros** *are macroscopic variables with minimal dynamical dependence over all optimisation runs at scale n*.

We consider the emergent *n*-macros across all candidate *spatial* scales and we define the *localisation perspective* of emergent dynamical structure as^3^.

##### Definition 4.

**Emergent dynamical structure** *is defined by the set of all emergent n-macros which together reveal the emergent dynamical structure across spatial scales of the whole system*.

This dynamical structure is revealed by identifying the degree to which the microlevel node contribution to emergent *n*-macros is distributed. This definition allows us to derive the *localisation* of *n*-macros in the original biophysical neural model.

In the context of our study, emergent macrovariables are defined as linear subspaces of the microscopic state space, where each dimension corresponds to a neural source or node in the network. The term localisation refers to the degree to which these macrovariables are aligned with specific axes of the microscopic space. Specifically, a macrovariable is considered *localised* when it aligns closely with one or a few of the original coordinate axes—meaning it has smaller subspace angles with these axes. This alignment indicates that the macrovariable primarily captures the dynamics of specific microlevel components, rather than being a distributed combination of many nodes. This concept of localisation is crucial for understanding how emergent dynamical structure forms within the network.

DI analysis applied to neurophysiological time-series will not necessarily reveal distinct emergent dynamical structure at the scale specified by the localisation of *n*-macros. Indeed, the present study develops a method able to characterise how functional integration and segregation relate to emergent dynamical structure across all spatial scales.

### Dynamical dependence minimisation in linear systems

We operationalise DI and the identification of emergent *n*-macros in simulated neurophysiological data using a linear approximation (see S1 Appendix: Granger causality, and S2 Appendix: Linear state-space modelling as to the appropriateness of the linear approximation). The microscopic state space is thus the Euclidean space 𝒳 = ℝ^*N*^, and coarse-grainings at spatial scale *n* are full-rank linear mappings (*n × N* matrices) *M* : ℝ^*N*^ → ℝ^*n*^.

For the coarse-grained macrovariable ***Y*** _*t*_ = *M* ***X***_*t*_, the dynamical dependence (Eq 1a) is invariant under nonsingular (invertible) linear transformations of the macroscopic state space 𝒴 = ℝ^*n*^. Thus we may consider two coarse-grainings *M, M* ^*′*^ to specify the *same macrovariable* if they are related by *M* ^*′*^ = Φ*M* for some nonsingular linear transformation Φ : ℝ^*n*^ → ℝ^*n*^. The space of all possible linear coarse-grainings at scale *n < N* —the space over which DD will be minimised—may consequently be identified with the set of *n*-dimensional subspaces of ℝ^*N*^. This defines a mathematical object known as a *Grassmannian manifold* [64, 65], *written 𝒢* _*N*_ (*n*). A coarse-graining *M* may be visualised as an *n*-dimensional subspace in the original microscopic state space ℝ^*N*^, on which the dynamics of the macroscopic variable ***Y*** _*t*_ = *M* ***X***_*t*_ play out. Intuitively, the coarse-graining map *M* projects the *N* -dimensional microscopic dynamics ***X*** down onto the macrovariable ***Y***, which resides on the associated *n*-dimensional subspace [30].

Grassmannian manifolds are a type of homogeneous Riemannian manifold; they are non-Euclidean spaces with a distinctive metric geometry and symmetries, on which we can do calculus. Given a microscopic process ***X***_*t*_, then, we can minimise the DD *T* (***X*** → *M* ***X***)—considered now as the objective function on 𝒢 _*N*_ (*n*) parametrised by the matrix *M* ^4^—using standard methods like gradient descent [64, 67] on the Grassmannian.

For the class of linear state-space models for the microscopic dynamics, moreover, both the DD and its gradient may be calculated explicitly; see S3 Appendix: Minimisation of dynamical dependence by gradient descent. In this study, we used gradient descent with an adaptive step-size [68] to minimise DD. Typically, minimisation is initiated at a uniform random *M* for a specific scale *n*, and allowed to run until it converges to a specified tolerance. However, as the optimisation landscape described by the DD is quite deceptive, with multiple local suboptima (minima), we repeat randomised optimisation runs many times to obtain acceptable minima for the specified scale.

### Characterisation the composition of linear macroscopic variables

A putatively emergent macroscopic variable is perhaps best thought of as a dimensionally-reduced subsystem of the global microscopic dynamics^5^. As explained above, in the linear case a macrovariable may be considered a projection of the microscopic process onto a subspace in the original microscopic Euclidean state space. In the context of modelling multi-region neurophysiological time-series data, we may get a sense of the extent to which different nodes (corresponding to neural signals in specific anatomical brain regions) in the brain network participate in, or contribute to, a macroscopic subsystem that we term an *n*-macro. We do this by invoking geometric intuition: consider, for example, a 2-dimensional plane in a 3-dimensional Euclidean space. The plane is uniquely identified if we know the *angles* between it and each of the three *x, y, z* coordinate axes in the 3-dimensional space (Fig 8(d)).

In the multi-channel recording scenario, microscopic coordinate axes correspond to recording channels (Fig 8(a-b)). Consider again the 3D example: if, say, the angle between axis *y* (corresponding to channel *y*) and the 2D plane associated with a given macroscopic variable at scale *n* = 2 is close to the maximum *π/*2, this tells us that neural activity in region *y* is *projected away* by the corresponding coarse-graining; region *y* does not participate strongly in the macroscopic process. As the angle approaches zero, participation is maximised.

With some caveats^6^, this generalises to arbitrary dimensions: to quantify participation of brain regions in a macroscopic subsystem we calculate the angles between each of the region axes and the macroscopic subspace; then *small* angles correspond to *high* contribution, and vice versa (see S5 Appendix: Principal angles & single-node contribution to macroscopic dynamics for details).

Subspace angles have another important use, as a metric to measure the similarity, or co-planarity between macrovariables. For example, when optimising dynamical dependence for (nearly-)DI *n*-macros, two gradient-descent runs may yield coarse-grainings *M*_1_ and *M*_2_, respectively, which we suspect may be identical, or nearly so. Measuring the angle between *M*_1_ and *M*_2_ can help us decide how similar they are: angles close to zero indicate high similarity. In the case where we have subspaces *M*_1_ and *M*_2_ of *different* dimensions *n*_1_ *< n*_2_, a subspace angle near zero indicates that *M*_1_ is nested in *M*_2_; that is, the *M*_1_ dynamics may be viewed as a self-contained subsystem of the *M*_2_ dynamics. Within the current study we only consider subspace angles from microlevel constituents to *n*-macros and leave the comparison across higher-order scales for future research.

#### Brain network model

Using The Virtual Brain (TVB) [9, 69], we simulate biophysical neural models by constructing a 5-node network. Each node’s local dynamics are governed by the Stefanescu-Jirsa 3D (SJ3D) neural mass model (NMM), a reduced model capturing the mean field activity of 150 excitatory and 50 inhibitory Hindmarsh-Rose neurons [9, 52, 53]. Spike-burst neurons are thought to be implicated as critical neural mechanisms underlying conscious processing [54, 55]. The SJ3D model, governed by six coupled differential equations (considered state variables), is detailed in S4 Appendix: Global brain network dynamics. The model parameters used have been optimised to fit resting-state EEG data, following [69].

We choose our microscopic level as represented by the SJ3D NMMs, and the macroscopic scales defined across higher-order spatial scales *n* = 2 and *n* = 3.

#### Local dynamics

The SJ3D neural mass model consists of an excitatory and an inhibitory neural mass, which are interconnected through the fast excitatory variable *ξ* and the fast inhibitory variable *α* (detailed in Table 2). These variables, along with the connectivity between the excitatory masses, define the dynamics at each node. The SJ3D model can generate various ensemble dynamics, including excitable regimes, oscillations, and transient spike-bursts [53]. The illustration below shows the construction of the NMM.

A key advantage of the SJ3D worth noting is its multi-modal construction. Modes represent different dynamical regimes in which the local dynamics can exhibit and represent distinct population dynamics which give rise to rich heterogeneous activity exhibited locally [9, 69, 70].

#### Global dynamics

Our simulations are based on a well-established evolution equation governing the dynamics of brain network models, adapted and modified from The Virtual Brain (TVB) for this study (details in Eq 27 in S4 Appendix: Global brain network dynamics). The structural connectivity, which defines the anatomical backbone of the network model, is represented by a weight matrix and a tract-length matrix. In the context of a whole-brain model (WBM), the weights matrix quantifies the strength of pairwise anatomical connections between brain regions, while the tract-length matrix measures the axonal fibre lengths between these regions.

The 5-node biophysical neural model used here is derived from an empirically-informed TVB structural connectome, which incorporates a biologically realistic tract-length matrix. This matrix is created through homotopical morphing, a computational technique that optimizes and aligns primate tracer imaging data with the human anatomical connectome, based on a Desikan-Killiany parcellation atlas [71]. To evaluate the plausibility of the emergent dynamical structure, constrained by network connectivity, we also perform the same analysis on an uncoupled structural connectome as a control condition. Throughout the experiments, the network connectivity was kept constant across both the coupled and uncoupled regimes, serving as a ground-truth model that constrains the dynamics revealed by varying degrees of functional integration and segregation (see Fig 2).

**Fig 1.**
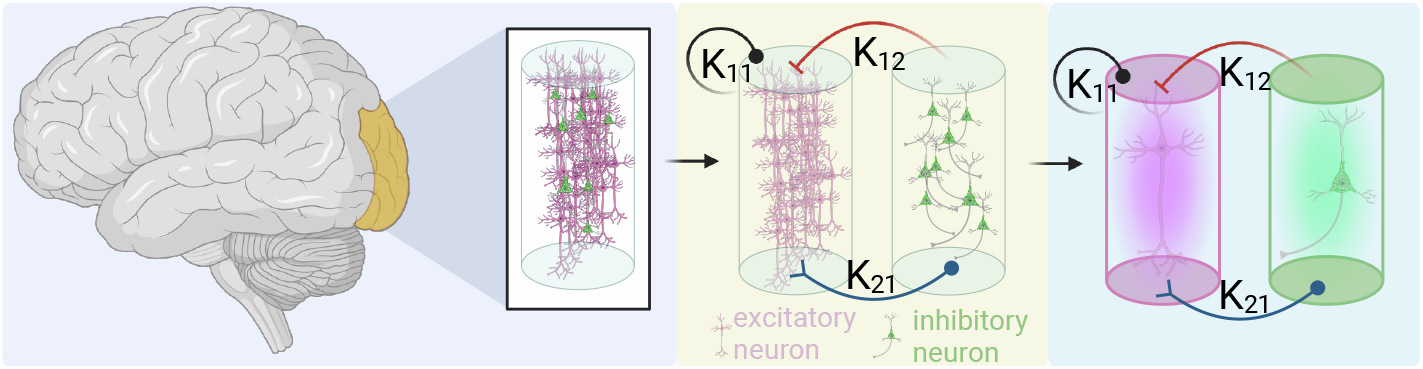
Network architecture of a Stefanescu-Jirsa model. *K*_12_ is the inhibitory-to-excitatory mass connectivity, *K*_21_ is the excitatory-to-inhibitory connection, and *K*_11_ is the excitatory-to-excitatory connectivity. The left-most panel indicates that each neural mass model, consisting of both, excitatory and inhibitory neural masses represents a single node in a region-based brain network simulation. Going from the central panel to the right-most indicates the reduced model of the mean-field approximations representing the ensemble activity on the excitatory (pink) and inhibitory (green) neural masses, respectively.

**Fig 2.**
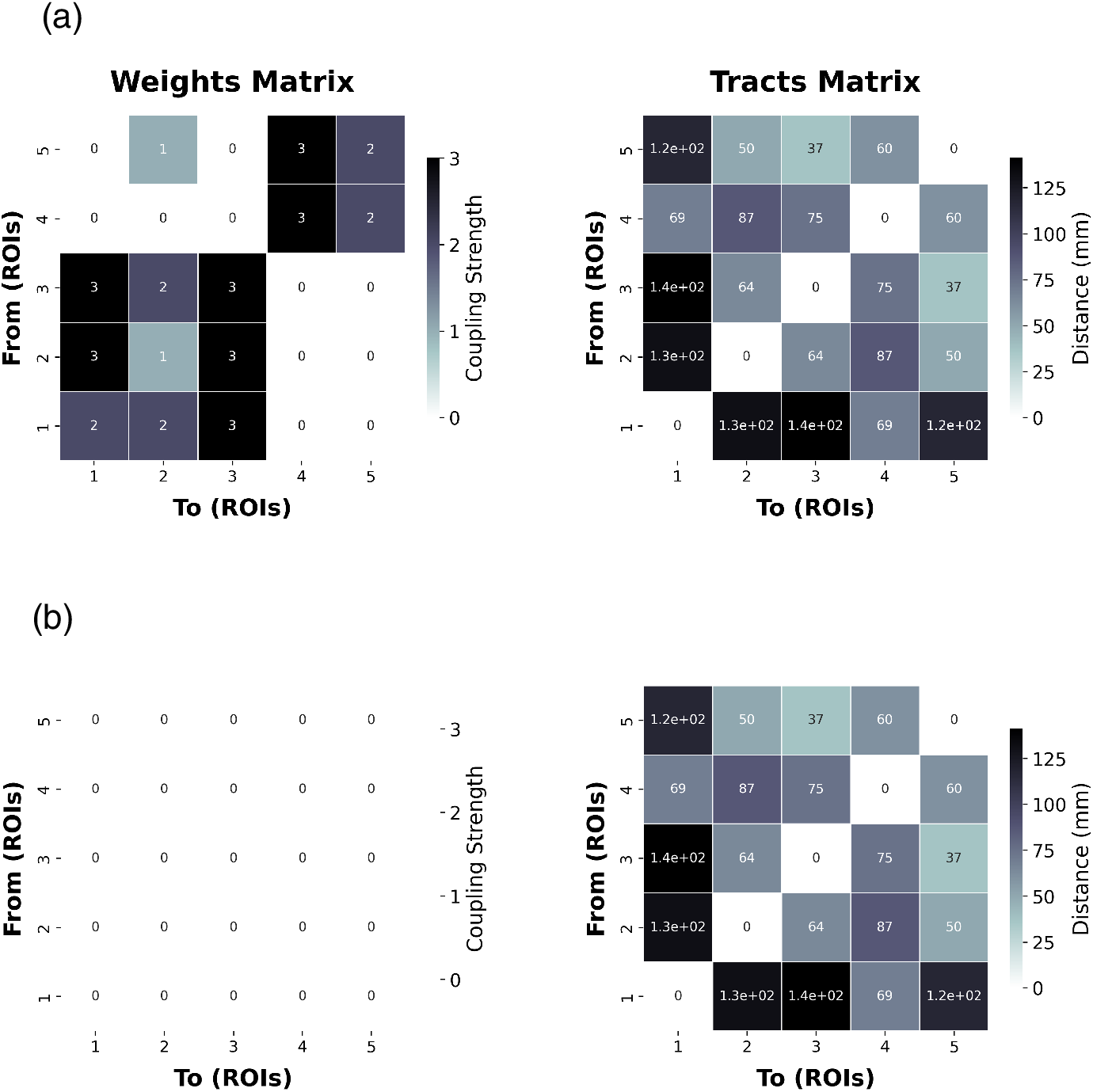
Structural connectivities defining the 5-node brain network models implemented in this study. *(a)* the coupled connectivity defined by the weights and tract-lengths matrices, and *(b)* the uncoupled connectivity.

To reiterate, two global parameters known to influence macroscopic brain dynamics [2, 57, 72] were manipulated: global coupling and dynamical noise. Functional integration is defined by the global coupling factor (*G* in Eq 27), which scales the influence of incoming activity from other nodes in the network. Conversely, dynamical noise, represented by the *v*_*i*_(*t*) term in Eq 27, reflects functional segregation by reducing the signal-to-noise ratio, thereby dampening the impact of external nodes on local dynamics.

#### Simulation protocol

A parameter sweep was performed on the values of global coupling (*G*) and dynamical noise (*η*) between 0.01 to 0.31 and 0.001 to 0.1 in 20 logarithmic steps, respectively (see Table 2 in S3 Appendix: Minimisation of dynamical dependence by gradient descent. To obtain the time-series data, we consider the *ξ* state variable at each node, representing the excitatory activity. The activity for each *ξ* was summed over the 3 dynamic modes [70] and then z-scored. The resulting time series represents the LFP-like excitatory activity of the respective brain region nodes [9, 53, 73]. For numerical stability a Heun stochastic integration scheme was used with a step size of *dt* = 2^−6^. This parameter can vary depending on the NMM used and the global parameter regime explored. For the present analysis it was identified that *dt* = 2^−6^ was consistently stable throughout the parameter sweep. Simulations were run for 5000 ms with the first 500 ms excluded to discount initial transients. Simulations were sampled at 256Hz to retain consistency with regularly deployed neurophysiological data acquisition methods such as electroencephalography (EEG) and intracranial-EEG (iEEG).

The resulting simulated neural time series for each of the 400 (20 × 20) simulations is subject to DI analysis using a linear approximations to estimate causal graphs and capture the emergent dynamical structure (see S1 Appendix: Granger causality for details).

## Results

To validate the capacity of our methodology to identify emergent macroscopic dynamics, we applied our DI analysis pipeline across 400 simulations with varying degrees of functional integration and segregation, using the network models defined in Fig 2. As mentioned, we leverage an uncoupled network model as a control condition, which is defined by the connectivity illustrated in Fig 2. For each simulation, we ran optimisations for a 2-macro and 3-macro in our 5-node biophysical network model. We selected a 5-node brain network to offer the simplest model in which we can vary the values of functional integration and segregation, while keeping the analysis and results as transparent as possible.

Fig 3 illustrates the minimal DD values of each emergent *n*-macro across simulations defined by varying global coupling and dynamical noise parameter values. Each matrix represents the entire bivariate parameter space, with dynamical noise varying along the *x*-axis and global coupling along the *y*-axis. The top row displays the DD values for the 2-macro and 3-macro in the coupled network, while the bottom row shows the corresponding results for the uncoupled network.

**Fig 3.**
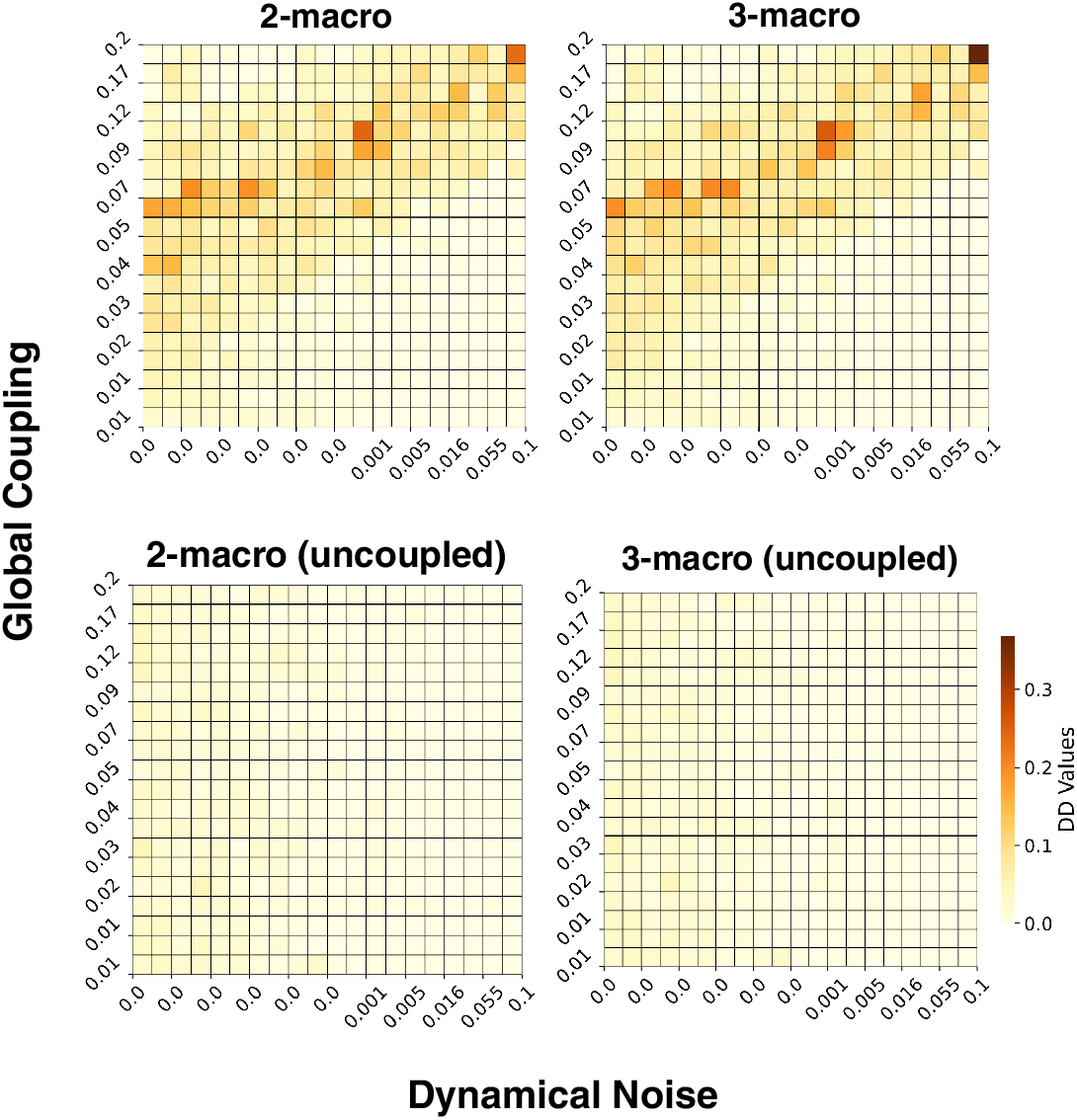
Minimal DD values of emergent *n* -macros in the 5-node network with local dynamics defined by a SJ3D NMM. Each matrix presents a parameter sweep across global coupling (y-axis) and dynamical noise (x-axis). The top row shows DD values for 2-macros and 3-macros in the coupled brain network. The bottom row presents the corresponding DD values for the uncoupled brain network. Darker colours indicate higher DD values, while lighter colours reflect lower DD values.

### Dynamical dependence peaks in a parameter regime balancing functional integration and segregation

Fig 3 illustrates that in the coupled network, there exists a parameter regime where the DD of emergent 2-macros and 3-macros is *higher*, indicating *lower* emergence of macroscopic dynamics—a pattern absent in the uncoupled network. Notably, this regime exhibits an consistent relationship where increases in both functional integration (global coupling) and functional segregation (dynamical noise) coincide. This balance results in the emergent dynamical structure being *maximised* through the localisation of contributions from specific microlevel nodes to the emergent *n*-macros, as dynamical noise increases proportionally to global coupling (see Fig 4 and 5).

**Fig 4.**
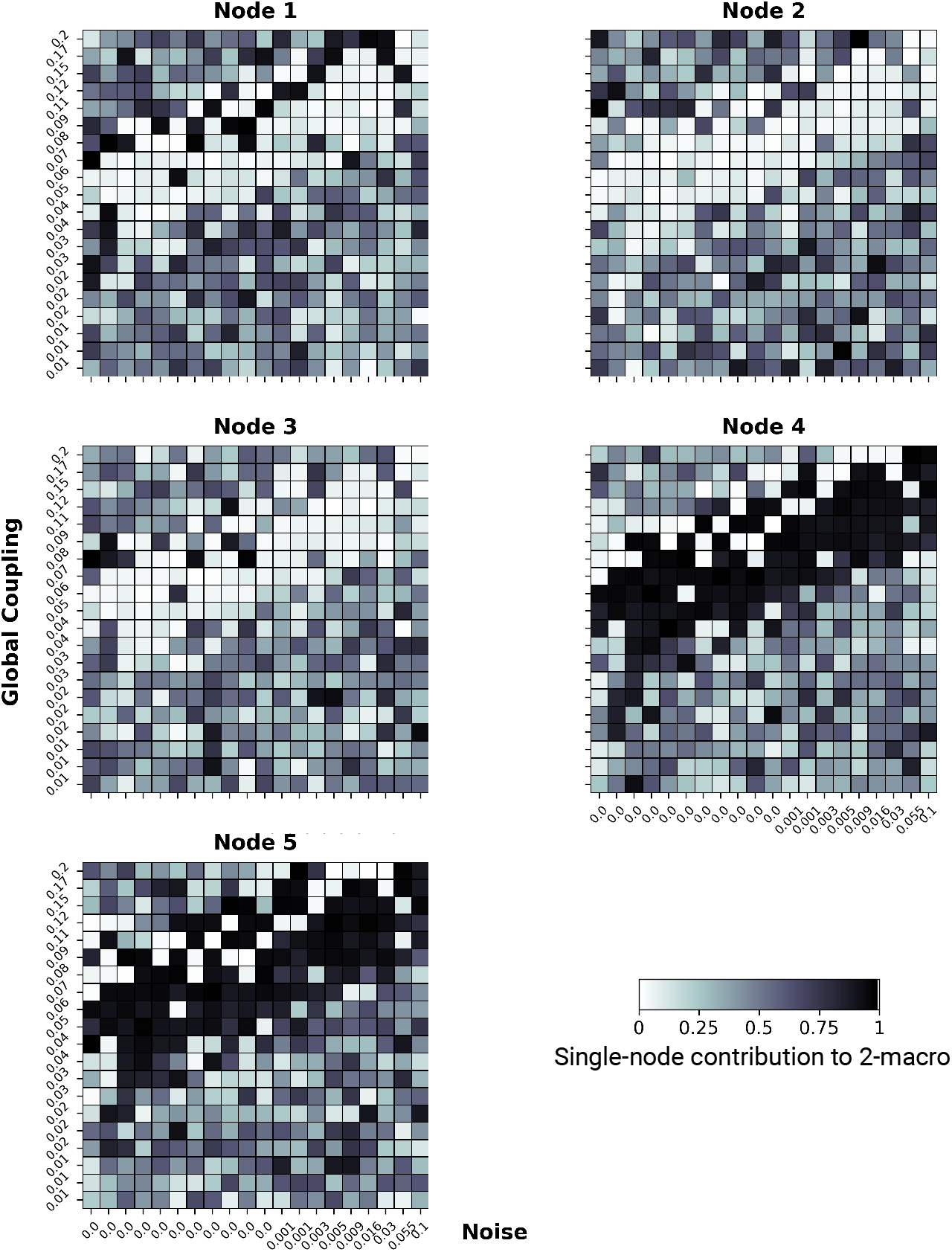
The single-node contribution to emergent 2-macros across simulations. Global coupling varies along the *y*-axis, and dynamical noise varies along the *x*-axis. Here, and on the subsequent figures identifying node contribution values on both axes are plotted on a logarithmic scale. Higher contributions are indicated by darker colours. The degree of localisation of single-node contributions within the emergent 2-macro is most distinct at a balance point between functional integration and segregation.

**Fig 5.**
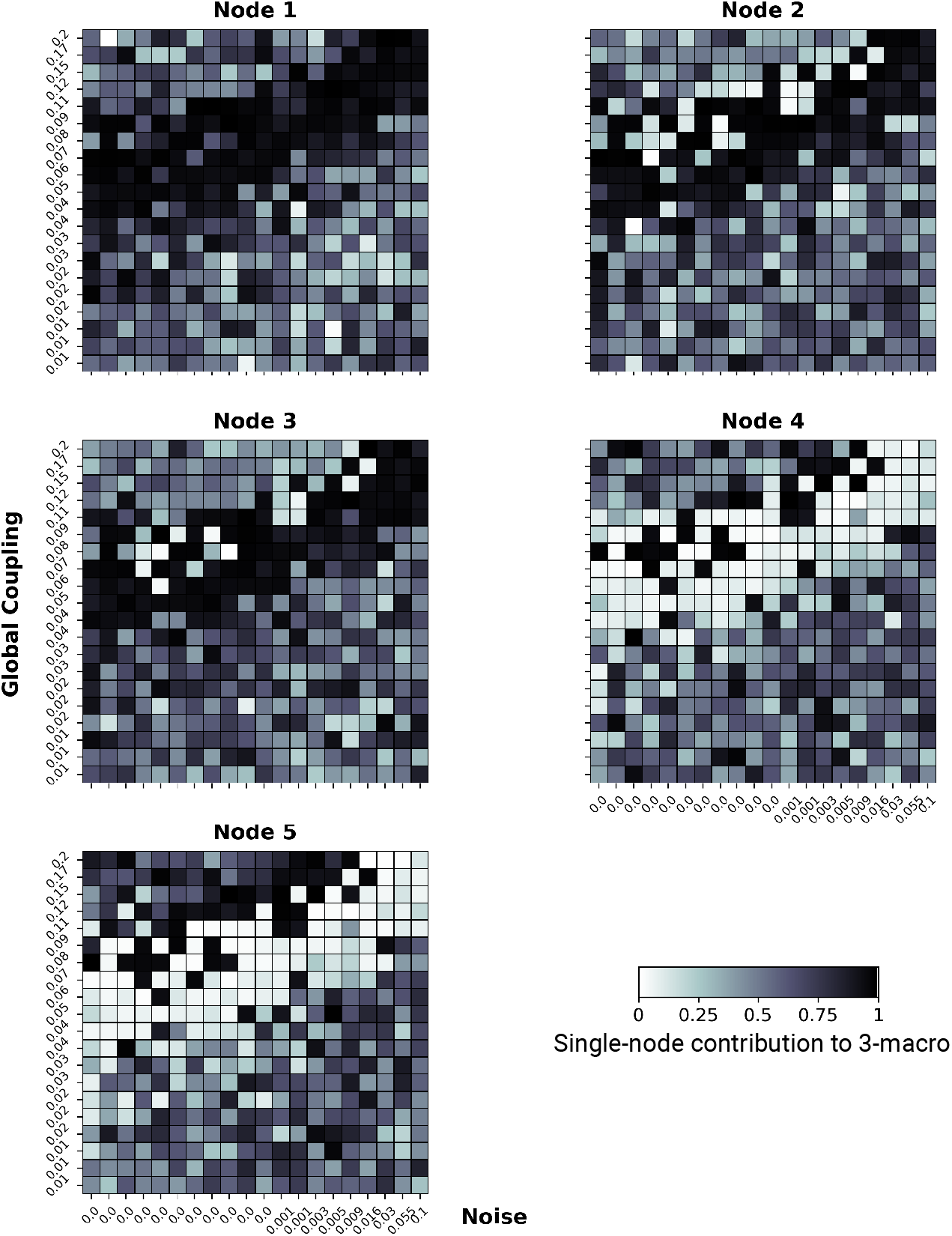
The single-node contribution to emergent 3-macros across simulations. Global coupling varies along the *y*-axis, and dynamical noise varies along the *x*-axis. Higher contributions are indicated by darker colours. The localisation of single-node contributions within the emergent 3-macro is evident at balanced points between functional integration and segregation, with a loss of localisation in extreme regimes.

This pattern suggests that the observed regularity across simulations is not merely a result of individual parameter values but rather emerges from the *interaction* between global coupling and dynamical noise. Their combined influence shapes the system’s dynamical structure, leading to higher DD (lower emergence) and maximised localisation of contributions at this balanced parameter regime. This finding underscores the importance of considering the interplay between functional integration and segregation when assessing both the emergence and the organisational structure of complex macroscopic dynamics.

### Identifying structure: Spatial localisation of single-node contributions in emergence *n* -macros across functional integration and segregation

We next performed a single-node analysis to examine the contributions of individual nodes to the emergent 2-macros across simulations. As shown in Fig 4 nodes 4 and 5 exhibit high contributions to the emergent 2-macro, while nodes 1, 2, and 3 show negligible contributions, particularly at the parameter regime where functional integration and segregation are balanced. This distinct organisation of node contributions towards an emergent 2-macro which is localised across 2 microlevel nodes is closely associated with the parameter regime characterised by higher DD (lower emergence) values (Fig 3). This indicates that at the balance point, the emergent dynamical structure is maximised through the localisation of contributions from specific nodes.

The degree of single-node contribution is represented as follows: a value of 0 (lighter colour) indicates that the node is not implicated in the emergent 2-macro, and a value of 1 (darker colour) indicates that the node is fully implicated. For a detailed explanation see S6 Appendix: Worked example.

Further analysis, as illustrated in Fig 4, reveals that in simulations where the parameter regime is dominated by either excessive functional integration or segregation, all nodes show varied degrees of contribution to the emergent 2-macro. This variability indicates a breakdown in the localisation of single-node contributions, which underpins the dynamical structure of the emergent *n*-macros. Notably the dynamical structure of the *n*-macros appears *randomly* distributive across the entire network.

By examining single-node contributions across all parameter regimes, we assess the degree of localisation within each *n*-macro, providing insight into the integrity of the emergent dynamical structure. Importantly, distinct localisation is most apparent in parameter regimes associated with higher dynamical dependence (lower emergence)—that is, at a balanced point of the co-existence of functional integration and segregation. In contrast, in regimes with extreme dynamical noise or global coupling, this localisation diminishes, leading to a more distributed contribution of nodes across the entire network and a subsequent weakening of the dynamical structure.

Similarly, Fig 5 shows that single-node contributions to an emergent 3-macro exhibit a distinct localised pattern at balanced points between functional integration and segregation. In this regime, nodes 1, 2, and 3 contribute significantly to the emergent 3-macro, while nodes 4 and 5 show minimal to no contribution, as reflected by the lighter colours in their respective matrices. However, when the parameter regime shifts towards extreme functional integration or segregation, this localisation of contributions weakens. The result is a more distributed contribution across all nodes, leading to a diminished and less coherent dynamical structure.

Finally, given that the uncoupled network serves as a control, we should expect the minimal DD-valued *n*-macros in such a system to reflect subspaces that span some combination of the original coordinate axes. That is, they are localised or distributed across some set of nodes. In fact, without the need for simulation, we can theoretically predict that for a macro dimension *n*, any subspace formed by linear combinations of *n* of the *N* original coordinate axes (microscopic state space) should exhibit close-to-zero DD values. Indeed, this expectation is supported by the results obtained in the lower graphs in Fig 3, where the 2- and 3-dimensional emergent macros identified by the optimisation process in the uncoupled system correspond to these close-to-zero DD subspaces.

Further, Fig 6 illustrates that in the uncoupled network, the contributions of microscopic nodes to the emergent 3-macros^7^ vary randomly across the entire parameter space, resulting in the absence of distinct localisation of the dynamical structure of emergent macros, even when functional integration and segregation are balanced. These findings suggest that the anatomical connectivity between nodes in the brain network is crucial in determining the localisation of dynamics and the contribution of nodes to emergent dynamical structure at the macroscopic scale. In the absence of coupling, the emergent dynamical structure is *not maximised*, and the DD values are close to zero, indicating higher emergence (lower DD) but without the distinct localisation that characterises the maximised emergent dynamical structure in the coupled network.

**Fig 6.**
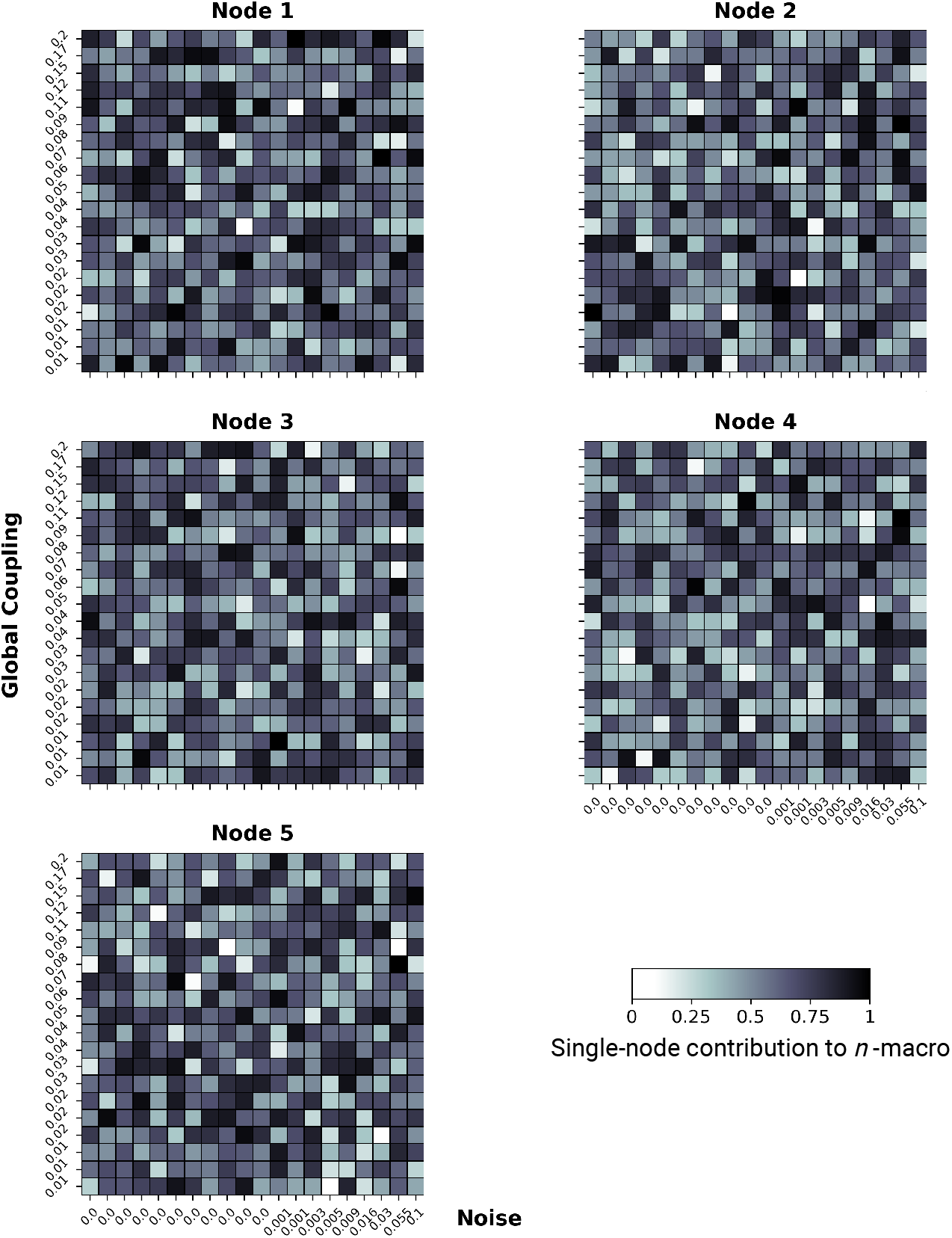
The single-node contribution to emergent 3-macros across simulations. Global coupling varies along the *y*-axis, and dynamical noise varies along the *x*-axis. Higher contributions are indicated by darker colours. The contributions of nodes to the emergent 3-macros vary randomly across the parameter space, indicating the absence of distinct localisation in the uncoupled network.

### Statistical significance of single-node contributions to emergent *n* -macros

Thus far, we have explored how and when macroscopic dynamical structure emerges through the localisation of single-node contributions to emergent *n*-macros. To assess whether these single-node contributions to emergent 2-macros and 3-macros across the parameter space are statistically significant compared to the uncoupled brain network (control condition), we performed a Wilcoxon rank-sum test. Specifically, this analysis compares the distribution of node weights across all simulations for the coupled and uncoupled regimes (Fig 4, 5, and 6). The statistical comparison is conducted across the entire parameter space explored, not just within the parameter regime where we qualitatively observe distinct localisation of the emergent macroscopic dynamical structure. By performing the statistical comparison across the full parameter space, we ensure a robust and comprehensive assessment of the significance of single-node contributions to emergent macroscopic dynamical structures, thereby avoiding potential biases that could arise from selectively focusing on regions where qualitative observations suggest distinct localisation.

Consulting Fig 7, the Wilcoxon rank-sum test reveals a significant difference in the contribution of node 4 to an emergent 2-macro in the coupled brain network compared to the uncoupled brain network used as a control condition (*Z* = 2.08, *p <* 0.05). No significant contribution was observed from any other nodes. Interestingly, despite the qualitative illustration in Fig 4 showing distinctly higher contributions from node 5 to the emergent 2-macro, this node did not exhibit statistical significance when compared to the control Fig 6. This discrepancy might be due to the statistical analysis being conducted across the entire parameter space, rather than being confined to the regime where distinct localisation is observed. However, it is curious that node 4 still shows significance under the same conditions, suggesting that the lack of significance for node 5 may not be fully explained by this alone.

**Fig 7.**
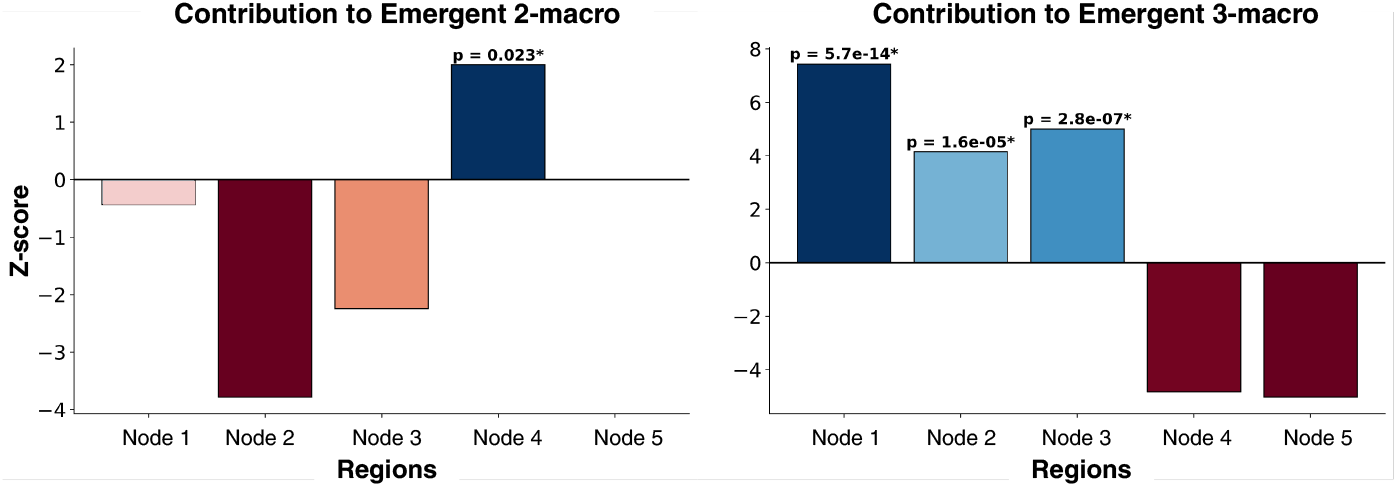
Wilcoxon rank sum test for single-node contribution. (*left*) Emergent 2-macro and (*right*) Emergent 3-macro, in the coupled network compared to the uncoupled network across a parameter sweep of global coupling and dynamical noise

**Fig 8.**
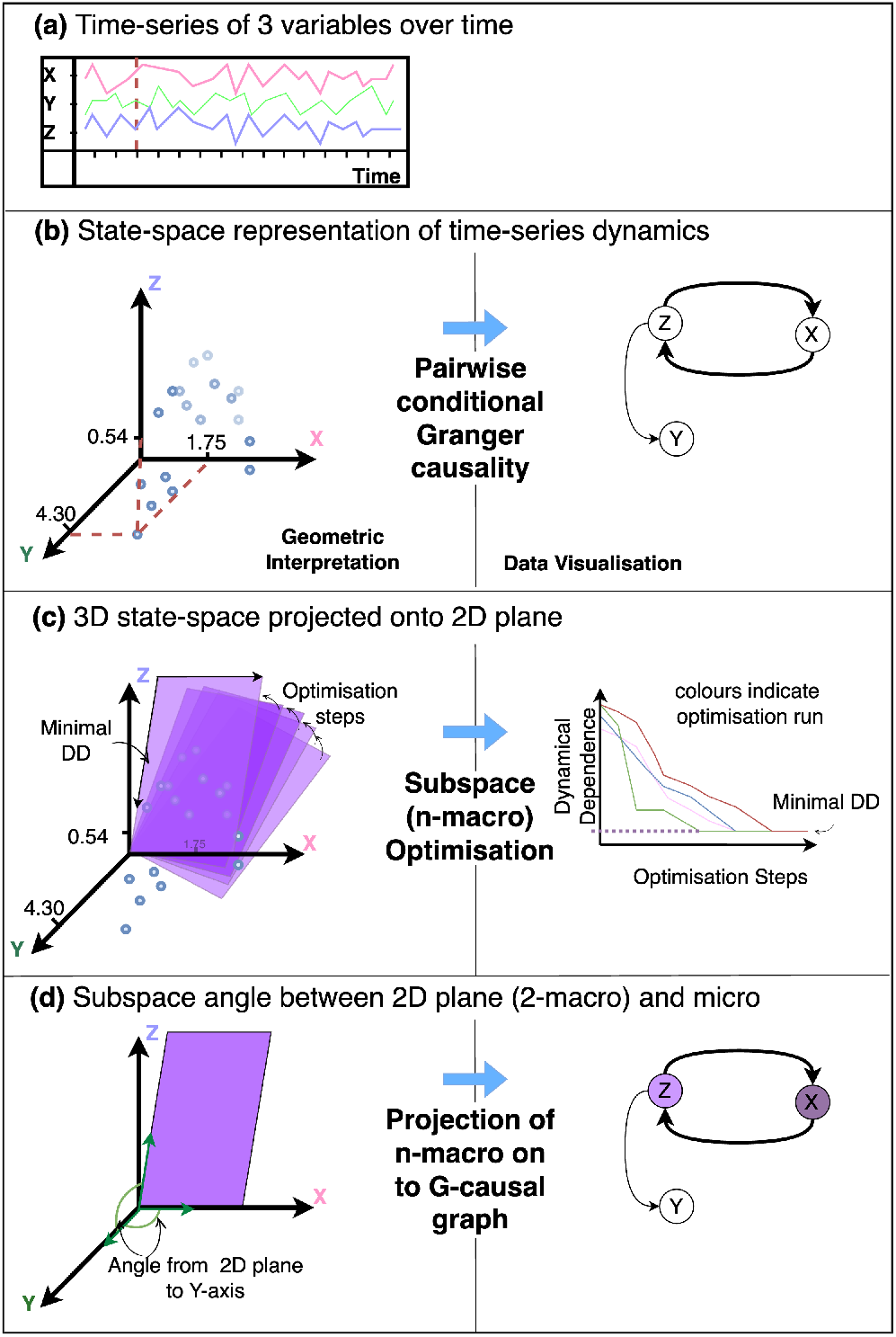
Geometric interpretation and data visualisation equivalent describing the DI algorithmic pipeline. *(a)* a time-series capturing the dynamics of 3 variables, brain regions, or electrodes. *(b)* a state-space model estimated from the time-series dynamics from which we calculate the pairwise-conditional Granger-causal statistics representing directed information flow. *(c)* A 2D plane embedded in the 3-dimensional system. This plane represents the lower-dimensional macroscopic variable – a 2-macro – capturing the projection of the 3D dynamics onto a 2D space (dimensional reduction). In optimisation, the plane shifts incrementally toward decreasing levels of dynamical dependence, moving closer to a local or global minimum. *(d)* The minimally dynamical dependent plane is selected as the emergent 2-macro. To determine the single-node contribution to the emergent 2-macro distances between the axes of the original 3D system, and the axes of our emergent 2-macro are computed. A projection of the emergent 2-macro back onto the G-causal graph results in a visualisation of the degree of contribution from from each node.

Similarly, a Wilcoxon rank-sum test reveals a significant difference in the contributions of nodes 1 (*Z* = 7.69, *p <* 0.0001), 2 (*Z* = 4.13, *p <* 0.0001), and 3 (*Z* = 4.98, *p <* 0.0001) to an emergent 3-macro in the coupled network compared to the uncoupled network. In contrast, nodes 4 and 5 do not show significant contributions.

## Discussion

Overall, our results reveal that when the co-existence functional integration and segregation are finely balanced, the dynamical dependence (DD) of macroscopic variables is *higher* compared to other parameter regimes. This indicates that macroscopic dynamics are less emergent and more dependent on the underlying micro-level dynamics at these points. However, the emergent dynamical structure—defined by the localisation of contributions to macroscopic variables—is *maximised* under these conditions, with specific micro-level nodes distinctly contributing to the macroscopic dynamics. Conversely, deviating from these balanced points leads to a *lower* DD (indicating more emergent macroscopic dynamics) but results in a loss of localisation, where the contributions of micro-level nodes become more distributed.

We developed and outlined a complete computational method for identifying emergent dynamical structure in neural models. By modulating global coupling—pushing the system toward functional integration—and dynamical noise—pushing the system toward functional segregation—we examined how their balanced coexistence influences both the dynamical dependence and the localisation of contributions to macroscopic variables in a biophysical neural model. This approach allowed us to uncover the nuanced relationship between minimised emergence (in terms of higher DD) and maximised emergent dynamical structure (in terms of localisation) at balanced integration and segregation.

First, in a worked example, we illustrated that even in simple toy systems, achieving absolute dynamical independence is nearly impossible, as predicted by theoretical claims [30], and that the weighting of each node’s contribution rarely equals zero It is important to note, however, that these node contributions primarily reflect the localisation of the *n*-macros within the original system’s state space, rather than directly indicating their emergent characteristics on the higher-order scales themselves. For instance, in a more general scenario where the network connectivity is arbitrarily rotated in a high-dimensional space, the emergent *n*-macros would remain unchanged (due to the transformation-invariance of the DI framework), but the node contributions could appear more dispersed and potentially random. This highlights the distinction between node contributions and the emergent properties on the higher-order scales. While this limitation means that node contributions alone may not fully capture the *higher-order* structure, they remain valuable for revealing how macroscopic dynamical are spatially localised within the network: an aspect of the *emergent dynamical structure* of the neural model. This localisation insight is crucial for understanding the aggregation of local interactions into global dynamics, complementing other methods that may better capture higher-order interactions. As mentioned a potential utility in capturing the structure of the higher-order interactions can be through similarity matrices (see S6 Appendix: Worked example, for a worked example on a simple system). By integrating single-node analysis with other methods, we can achieve a more comprehensive understanding of the emergent dynamical structure in empirical data.

Next we constructed neural models that were intentionally designed with modular structures to assess how functional integration and segregation effect the emergent dynamical structure even with ground truth modularity. While this setup aids in illustrating the concept, it may not fully represent real-world neural data where *n*-macros do not necessarily correspond to such straightforward structure via analysis of their spatial localisation.

This is important, because in our approach coarse-graining within the DI framework is not formalised as a partitioning function that simplifies a complex high-dimensional network into a more manageable structure, as seen in other formal approaches to emergence [74]. Instead, it serves as an information-theoretic dimensionality reduction technique that captures lower-dimensional descriptions of whole-system dynamics across spatial scales. Because it does not rely on strict partitioning, this approach is well-suited to address the graded nature of contributions from individual nodes within the network. By avoiding the rigid boundaries imposed by partitioning, our method allows for a nuanced analysis of how these contributions vary, enabling us to assess whether the dynamical structure is more localised—where a few nodes dominate—or more distributive—where contributions are spread across many nodes—as the parameter values change. This flexibility is crucial in capturing the complex interplay between functional integration and segregation, and in understanding how local interactions aggregate into global dynamics. Importantly, this method also differs from other dimensionality reduction techniques, such as PCA, t-SNE, and UMAP [75], as it focuses on capturing lower-dimensional descriptions of dynamics derived from the interactions of microlevel constituents, rather than merely accounting for variance or relying on algorithmic clustering. Following this, we now discuss some key results.

### Localisation of emergent *n* -macros: defining the emergent dynamical structure

Our results demonstrate that when the co-existence of functional integration and segregation are finely balanced, the emergent macroscopic dynamical structure is *maximised* through the localisation of single-node contributions to the emergent *n*-macros. Specifically, at these balanced points, the whole-system dynamics exhibit distinct patterns of single-node contributions to the emergent *n*-macros, rather than random or evenly distributed patterns. This maximised localisation defines the emergent dynamical structure at the balanced points of integration and segregation.

This distinct structure is evident from Fig 4 and Fig 5, where nodes 1, 2, and 3 show negligible contributions to the emergent 2-macro, while nodes 4 and 5 contribute significantly to the emergent 2-macro at the balanced points. Conversely, for the emergent 3-macro, nodes 4 and 5 show negligible contributions, while nodes 1, 2, and 3 contribute significantly. Statistically, we do not have a definitive explanation for why node 5 does not show significant contributions in certain cases while node 4 does (see Fig 7). We suspect that there might be a hidden bias in the optimisation procedure used for the DI analysis, particularly in the case of the uncoupled network. In theory, the optimisation should identify 2-macros that are combinations of pairs of coordinate axes uniformly across all pairs. However, it is possible that the procedure is biased towards certain pairs more than others. This issue requires further investigation into the optimisation procedure across the coupled and uncoupled networks to fully understand the underlying reasons.Moreover, when moving away from the balanced points—either by increasing functional integration or segregation—the lack of distinct, localized single-node contributions to emergent *n*-macros leads to varied, distributed contributions from all micro-level nodes. This distributed nature induces a loss of distinct dynamical structure over the micro-level nodes that is otherwise observed at the balanced points. Interestingly, this loss of distinct structure is accompanied by relatively *lower* values of dynamical dependence (DD) for emergent *n*-macros, suggesting *higher* emergence of macroscopic dynamics.

While it might initially seem counterintuitive that a distinct, localised emergent dynamical structure occurs alongside *lower emergence* of macroscopic dynamics (as indicated by higher DD values), this observation remains consistent with the DI framework. At the balanced points, the DD is higher, indicating that the macroscopic dynamics are less emergent and more dependent on the underlying micro-level dynamics. The distinct structure does not necessarily imply that the emergent *n*-macros should possess a greater degree of dynamical (informational) closure. Crucially, the dynamical closure of the emergent *n*-macros is determined in relation to the microscopic processes alone, and not in comparison to other *n*-macros discovered across spatial scales.

Furthermore, lower dynamical dependence (higher emergence) does not necessarily mean that the emergent *n*-macro predicts itself well, i.e., that it is self-determining or autonomous (see [30]). Rather, it indicates that the macroscopic dynamics are more independent from the microscopic base. Through the same optimisation procedure, one could, in principle, reveal an entirely Gaussian, white-noise process as an *n*-macro that is independent of the micro-level constituents. Consequently, these ‘noisy’ macroscopic variables might not be dynamically relevant for the whole-system dynamical structure. However, while such a macroscopic variable might initially seem of limited interest, identifying these white-noise macros could be useful depending on the empirical question. They can potentially be factored out, allowing researchers to focus on the core, non-trivial dynamical structures that are more informative. From the results obtained, we suggest that the lower dynamical dependence (higher emergence) values observed when increasing dynamical noise (mediating functional segregation) could be driven by the emergence of such noise-dominated macros.

Consequently, we speculate that emergent macroscopic variables accompanied by relatively higher dynamical dependence (lower emergence) could be expected at the balanced points. This suggests that for the emergent *n*-macros to hold any dynamical relevance within the whole-system dynamics, some degree of dependence between the macroscopic process and the microscopic process may be necessary. However, the actual relevance of the emergent *n*-macros identified by DI analysis is ultimately an empirical question. It is crucial to carefully assess the functionality and significance of the *n*-macros, recognising that not all identified macros may contribute meaningfully to the overall system dynamics.

Lastly, Fig 6 indicates that the parameter regime associated with the emergence of macroscopic dynamical structure is absent in an uncoupled network. This finding underscores the importance of interactions between micro-level constituents in driving the emergent macroscopic patterns of activity in neural models. Additionally, we demonstrated that statistical testing can effectively discern single-node contributions to emergent *n*-macros across different conditions, providing a rigorous method for determining *n*-macro localisability in the original state space. In this study, we focused on the contribution levels of each node to the dynamics of the emergent *n*-macros, comparing these contributions with those in the uncoupled network using a Wilcoxon rank-sum test.

### Beyond indices: Uncovering the dynamical structure of complexity and emergence

Our approach could challenge existing intuitions about organisational complexity in both computational [76, 77] and biological [78, 79] systems. Furthermore, it explores the association between organisational structure and the optimal balance between integration and segregation [59, 80, 81]. Our methodology uniquely provides a robust operational approach to identify the dominant dynamical structures underlying global brain states, which are often indexed by measures of criticality [6, 21, 82] and neural complexity [59, 83–86].

An attempt to clarify the relationship between the integration-segregation balance and criticality has been attempted [25]. In general, criticality is commonly associated with systems at a phase transition, often characterised by power-law dynamics [87]. Theoretically, criticality refers to the point at which a system transitions between different phases, typically marked by scale-free properties [6]. However, the term”criticality” can be somewhat ambiguous, as it is empirically measured in various ways [6, 82], often alongside measures of complexity or signal diversity [88].

In this context, both measures of criticality and complexity serve as indices that refer to underlying organisational structure within the system. For instance, the power-law structure, such as the 1/f curve, indicates that the system exhibits scale-free behaviour, where smaller clusters of activity are more common than larger ones. This reflects a form of organisational complexity that is indexed by the criticality measure. Similarly, measures of neural complexity aim to index the degree of organisational structure within brain dynamics.

While remaining agnostic to specific measures of criticality or complexity, our work seeks to vary the two parameters that are believed to influence both, with the aim of providing a complementary method that goes beyond empirical indices to identify the underlying dynamical structure and its level of emergence. An exciting avenue for future research will be to compare our method with other neural complexity measures—which actually peak during a balanced point of integration and segregation—to explore how these measures deviate from each other. Characterising the emergent dynamical structure of global brain states in relation to these indices, using both observational and perturbed datasets, could offer significant advancements in our understanding of complexity in neuroscience.

Building on existing studies that explore the macrostates of brain activity [31, 37, 39, 89, 90], as well as those quantifying synergy between brain region pairs [49, 51, 91], our approach offers a complementary perspective by uncovering the macroscopic dynamical structure derived from microscopic interactions and quantifying its dependence on these interactions. Unlike other views of emergence, such as synergy, which captures a specific type of higher-order interactions between brain regions, DI provides a framework for understanding dynamical closure at the macroscopic level. Thereby identifying lower-dimensional subspaces that might serve as independent communication subspaces for regions. This approach not only reveals emergent macroscopic variables across spatial scales but also highlights the system-wide organisation that arises from the underlying dynamics. While existing methods applicable to whole-brain modelling might effectively capture macroscopic activity, they often lack a concept of emergence. Integrating these tools to form a comprehensive understanding of emergence in the brain represents an intriguing direction for future research. Further research will need to expand the application of our framework to larger artificial networks or neurophysiological data.

### Bridging emergence, coarse-graining, dynamical closure, and dimensionality reduction

This work contributes to the broader effort to (i) quantify and detect coarse-grained macroscopic dynamics, and (ii) provide a precise methodology for further application to large-scale neurophysiological data. In particular, we consider the relation of our approach to effective information-based measures of causal emergence [46, 74], informational closure [62, 63, 92], and common dimensionality reduction techniques [66, 75]

First, we examine the distinction between coarse-graining in the DI framework and coarse-graining within the context of effective information-based causal emergence [46, 74, 93, 94]. In causal emergence, coarse-graining involves recasting a complex network into non-overlapping partitions using a hard-partitioning function and evaluating the effective information of the resulting higher-order network. Effective information is measured by balancing the average degeneracy and determinism within the network. A partitioned graph that exhibits higher effective information than the original is considered causally emergent. This approach can be applied even without interventionist methods of causality [95, 96], extending across various bidirectional networks [74, 97].

In contrast, coarse-graining within the DI framework focuses on the degree of informational or causal closure of a macroscopic variable, aligning more closely with the principles of statistical mechanics. Instead of partitioning the network into sub-networks, this approach involves partitioning the microscopic *state-space* (which partitions the dynamics, not the nodes) into subsets *f* ^−1^(*Y*), where *Y* represents the macroscopic states. This method captures how individual microlevel nodes contribute to these macroscopic variables, which is much closer aligned to statistical mechanics descriptions macroscopic (ensemble) properties emerging from microscopic interactions [8, 98]. DI serves as a dimensionality-reduction technique that reveals low-dimensional dynamics as self-contained systems, without transforming the network into hierarchical structures. Given its ability to capture the distributive nature of macroscopic variables and their degree of dependence on the microlevel, DI aligns more closely with a heterarchical perspective of dynamical structure [99].

In support of a dynamical closure perspective on the emergence of macroscopic dynamics in the brain, it is important to recognise that a core principle of brain organisation is its function as a highly distributed information system [100–102], where local (microlevel) functional units integrate to generate macrolevel dynamics. These dynamics are not fixed but fluctuate over time, involving the transient recruitment of various microlevel regions [102]. This suggests that the brain’s global dynamics are driven by overlapping compositions of regions, without clear distinction between the microlevel parts and their relation to the whole. Regions within the brain are dynamically organised in response to functional and computational demands, reflecting the adaptive and non-static nature of brain dynamics across spatial scales. Although we worked on a 5-node simulated network in which these challenges are not present, our approach can adapt to the challenges of larger or real brain networks because it emphasises dynamical closure, focuses on the emergent dynamical structure, and examines to what degree microlevel nodes contribute to macroscopic dynamics—allowing for us to assess the degree of localised or distributive activity.

DI distinguishes itself from other formal approaches to informational closure, such as those by [103] and [63], by extending the concept beyond absolute informational closure (perfect dynamical independence) and accommodating non-Markovian dynamics. This allows DI to capture the nuanced interdependence between macroscopic and microscopic processes.

While [92] explore informational closure as a framework for conscious processing, our approach remains neutral on such interpretations. Additionally, unlike Chang and colleagues’ consideration of direct information flow between macroscopic and environmental variables, our framework imposes a supervenience condition (See Eq 2), ensuring that no new information emerges at the macroscopic level beyond what is determined by the microscopic processes. This means that the microscopic fully dictates the information shared with the macroscopic dynamics.

Finally, although DI results in dimensionality reduction, it offers distinct advantages over commonly used techniques such as Principal Components Analysis (PCA) and t-distributed Stochastic Neighbour Embedding (t-SNE). PCA identifies components of maximum variance, and t-SNE preserves local relationships between data points [66, 75, 104]. However, neither method explicitly accounts for the time-dependence of neural activity to capture low-dimensional dynamics. In contrast, DI operates as an information-theoretic dimensionality reduction technique, directly mediated by microlevel interactions by the temporal structure of the time series.

### Limitations and open questions

DI, in it’s current framework, is based on Granger causality. Although Granger causality is well-defined for non-stationary processes, it is notoriously difficult to estimate in these cases, and as such is not considered here. Thus we assume the wide-sense stationarity of neurophysiological data, and might not be able to capture physiologically-relevant non-stationarities in the time-series strongly correlated with specific brain states. An example can be illustrated by brief neural oscillations (e.g., sleep spindles), or waves (e.g., K-complexes, or epileptic activity) as well as transient responses to external or internal stimuli. However, linear approximations have been argued to optimally capture macroscopic neural dynamics [105], particularly in resting-state activity.

Furthermore, fitting a VAR or SS model for G-causality estimation assumes a linear model rather than a linear process, which is a subtle but important distinction. While the model assumes linearity in its structure, this does not necessarily mean that the underlying time series must be linear. The presence of non-linearities in the data does not inherently imply that the fitted model will be unstable (see [44] for an in-depth discussion); rather, it suggests that the model may not fully capture the effects of those non-linearities. In fact, if the SS or VAR model is stable, then Granger causality inference and DI analysis remain well-defined.

A more precise understanding is that the model, if stable, induces a linear representation of the underlying data. The key issue then becomes whether this induced linear model can sufficiently account for the non-linearities present in the time series. Wold’s decomposition theorem [106] guarantees that any stationary process can be decomposed into a linear model, although this model may require an infinite order, making it potentially non-parsimonious for time series generated by a nonlinear process [107]. Thus, the impact of nonlinearity on G-causality estimation is nuanced and complex. The critical question remains whether the linear model, when applied to nonlinear data, provides a sufficiently accurate representation of the underlying dynamics to make valid inferences. Though preliminary research suggests the advantages of linear models [105], this remains an interesting and open area for future research.

One limitation of our study is that the neural networks we employed are not functionally specialised—they are not designed to perform specific tasks or processes. In biological brain networks, functional specialisation is a fundamental characteristic that influences how integration and segregation manifest in neural dynamics. Therefore, it is possible that functionally specialised systems might exhibit different patterns of dynamical dependence and emergent dynamical structures. To address this, future research could explore simulations of networks with functional specialisation, perhaps by incorporating task-specific modules or connectivity patterns. Testing our methods in such simulated environments would help determine whether the observed results generalise to systems that more closely resemble the functional organisation of the brain.

While our study employs a relatively small 5-node neural model, which allows for detailed exploration of emergent *n*-macros, we acknowledge the importance of applying these methods to larger, functionally specialised systems. In ongoing work, we have begun to extend our techniques to whole-brain EEG data, demonstrating that our approach can be scaled up to analyse complex neural dynamics at the whole-brain level. This progression not only addresses the computational challenges associated with larger models but also brings us closer to understanding emergent dynamical structures in more realistic neural systems.

Ultimately, the dynamical relevance and implications of emergent *n*-macros for the systems under study remain an open empirical question. It is plausible to consider that emergent *n*-macros with many equally implicated contributions from microlevel nodes could function as low-dimensional communication subspaces through which higher-order interactions are mediated [108]. These *n*-macros, by capturing distributive contributions across the network, may represent the channels through which complex, coordinated dynamics occur at a macroscopic level. With DI we can establish the degree of localisability or distribution of these communication subspaces (*n*-macros). This concept of communication subspaces presents a compelling direction for future research.

## Conclusion

This work provides methodological, empirical and theoretical contributions to the exploration of emergent dynamics in complex neural systems. From a methodological and empirical perspective, we demonstrate that the balance between functional integration and segregation significantly influences the emergent dynamical structure across macroscopic spatial scales in biophysical neural models. Our results revealed that a distinct dynamical structure is identified by the spatial localisation of *n*-macros at a balanced point between integration and segregation, where specific microlevel nodes predominantly contribute to specific emergent *n*-macros. In contrast, this organised structure becomes less localised and more distributed in parameter regimes marked by either excessive integration or segregation. These results illustrate the heuristic power our approach may have when applied to biological systems, in identifying the brain structures participating in emergent dynamics.

From a theoretical perspective, this work contributes to the broader agenda of moving beyond indexical measures of complexity and emergence to identifying the underlying structure of dynamics that underpin quantities like these. Progress in identifying dominant emergent macroscopic patterns has implications for defining stable global brain states and understanding how the brain organises itself to meet the computational demands of whole-brain function. The findings here represent a promising step towards leveraging DI for empirical investigations into dynamical properties that go beyond traditional complexity indices, offering a more qualitative perspective of brain organisation.

## Glossary

Emergence: The phenomenon where higher-level patterns or behaviours arise from interactions of lower level constituent units.
Emergent Dynamical Structure: Macroscopic *patterns* of activity across spatial scales that emerge from the interactions between microscopic variables in a system.
Integration: The tendency of different parts of a system to work together to form a cohesive whole, often through coupling between units.
Segregation: The tendency of parts of a system to behave independently, reducing the interactions and coupling between them.
Surjective: A function that maps every element in the target set (output) to at least one element in the domain (input).
Stochastic Process: A process that involves randomness, where the next state is not fully determined by a set of previous states in it’s past.
Coarse-Graining: A technique that reduces the dimensionality of a system by grouping together similar states or variables into a single, larger-scale description.
Macroscopic Variable: A higher-level, large-scale variable that describes the collective behaviour of many microscopic variables.
Microscopic Variable: A small-scale variable that represents the state or behavior of a component in a system.
Dynamical Independence (DI): A measure of how independent the dynamics of a macroscopic variable is from the underlying microscopic variables.
State-Space: The set of all possible states that a system can occupy.
Partition: A division of the state-space into distinct, non-overlapping subsets.
Parameter Sweep: The process of systematically varying parameters in a model to observe their effects on the system’s behaviour.
Distributed / Local: Refers to whether a macroscopic process is spread across many variables (distributed) or confined to a few variables (local).
Neural Mass Model (NMM): A simplified mathematical model that represents the collective dynamics of neural populations, often used to describe brain activity.
Biophysical Model: A computational model that incorporates biological realistic details, capturing the physical and biological mechanisms underlying a system, such as neural dynamics.
Evolution Equation: A mathematical equation that describes how a system’s state changes over time, often used in neural models to represent dynamic processes.
Higher-order Scales: Larger-scale, macroscopic descriptions—across spatial scale—of a system that capture the emergent dynamics arising from interactions at the microscopic or lower-level scales.

## Supporting information

### S1 Appendix Granger causality

Estimating transfer entropy comes with some key challenges [61]. As a conditional mutual information, the state-of-the-art estimator uses the Kraskov–Stogbauer–Grassberger (KSG) nearest-neighbour algorithm [109–111], though kernel methods are also popular [110]. Due to inherent limitations of nearest-neighbour methods, the KSG estimator tends to overestimate TE, resulting in positively biased estimates (see Bossomaier *et al*. [61] for an in depth exposition). Furthermore, in lieu of known analytic sampling distributions, subsampling techniques must be employed for statistical testing, which may not systematically mitigate the bias inherent in nonzero TE estimates [112, 113]. Another issue with nonparametric estimators such as KSG, is that due to a curse of dimensionality which relates to the number of time-lags used in both source and target variables for TE estimation. In practice, one can restrict to just a few time lags, but this is known to risk skewing TE estimates [114]. Because of the inherent computational inefficiency and lack of a known analytic sampling distribution for statistical inference we leverage the close relationship between Transfer entropy and Granger causality [115, 116]. We remark that, unlike transfer entropy, Granger causality may be decomposed in the spectral domain; we don’t however make use of spectral Granger causality in this study. Transfer entropy and Granger causality are equivalent under the assumption that the stochastic processes involved are jointly Gaussian [115]^8^; the equivalence extends to a broader class of exponential distributions [117]. More generally, for arbitrary (possibly nonlinear) Markovian predictive models, if the GC statistic is construed as a log-likelihood ratio (see below), then GC is asymptotically equivalent to TE in the large-sample limit [116]. Granger causality may be viewed, then, as an approximation to transfer entropy, and inherits the interpretation of the latter as *information flow*. In comparison to transfer entropy, Granger causality is generally more efficient to estimate, is able to accommodate longer histories, and common estimators have smaller bias and variance, as well as known analytic sampling distributions.

Granger causality reframes transfer entropy in terms of least-squares optimal prediction, as opposed to (conditional) mutual information. Suppose given a stationary, multivariate, continuous-valued, zero-mean stochastic process ***U*** (the *universe of information* [118]) specified by 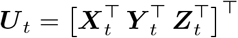, which is partitioned into three sub-processes ***X, Y*** and ***Z***. We say that the source variable ***X*** Granger-causes the target variable ***Y*** conditional on ***Z*** if the prediction of ***Y*** based on its own past and the past of ***Z*** is enhanced by the addition of the past of ***X*** to the predictor set, where prediction is in the least-squares sense, and (following [119]) quantified by the log-generalised variance^9^ [120] of the residual prediction error^10^.

More precisely, the *full* optimal least-squares prediction of ***Y*** _*t*_, based on the joint past ***U*** _*t*−*τ*_ of all variables^11^, is given by the conditional expectation

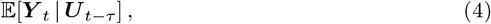

with residual error

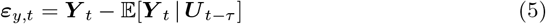

(the *y* subscript indicates that the target (predicted) variable is ***Y***). We use the notation *t* − *τ*, where *τ* is not a single value but represents a set of lagged time points, *τ* = 1, 2, 3, …, *n*, referring to the history of the process. In the case of an infinite history *τ* = 1, 2, 3, …, this set would extend indefinitely.

The *restricted* (or *reduced* prediction of ***Y*** _*t*_ is based on the joint past of the restricted process ***U*** ^R^ given by 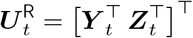, where the source variable ***X*** is omitted (we generally denote restricted quantities by a superscript “R”):

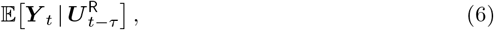

with residual error

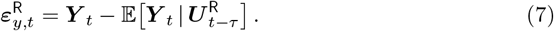

The full and restricted prediction error covariance matrices are respectively

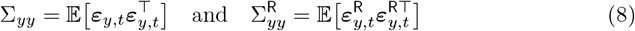

and, following [119], the Granger causality from ***X*** to ***Y***, quantifying the gain in prediction efficacy by inclusion of the source history ***X***_*t*−1 : −∞_ in the predictor set, is then defined as the log-ratio of generalised variances

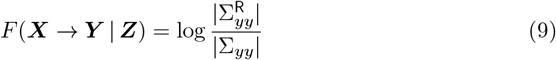

If there are no conditioning variables, then the above goes through with ***Z*** omitted.

The most common application of Granger causality to neurophysiological data is derivation of the *(Granger-)causal graph*. Given a multivariate (vector) process ***X*** with 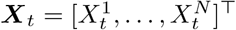, where the univariate processes *X*^*i*^ commonly represent single channels or regions of recorded data associated with specific anatomical brain regions, this is the directed, weighted graph of pairwise-conditional Granger causalities, defined as

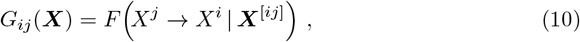

where the [*ij*] superscript indicates that the variables *X*^*i*^ and *X*^*j*^ are omitted from the full process ***X***. Thus *G*_*ij*_(***X***) represents the directed GC from channel *j* to channel *i*, controlling for indirect influences from other channels. The causal graph is often taken to summarise directed functional connectivity for neural systems.

In sample, the optimal predictions (conditional expectations) (4), (6) may be estimated from the data by standard Ordinary Least Squares (OLS), or another maximum-likelihood estimator, and if ***U*** is a Gaussian process, then the sample statistic corresponding to (9) is a log-likelihood ratio (*cf*. [116]); as such, under the null hypothesis of vanishing GC it has an asymptotic 𝒳 ^2^ distribution^12^.

The above exposition suggests that in sample both full and restricted predictions must be estimated separately; in practice, this is equivalent to estimating (full and restricted) vector-autoregressive (VAR) models. There is also a *single-regression* VAR estimator, where the restricted residual error distribution is derived analytically from the full model estimate. This estimator has smaller bias and variance than the conventional dual-regression estimator, and the null sampling distribution is known, at least in the unconditional case, to be a generalised 𝒳^2^ [121]; see [122].

Alternative routes to estimation of Granger causality beyond VAR modelling exist: the first is a parametric approach based on linear state-space modelling, while the second, nonparametric method is based on *spectral factorisation* of the cross-power spectral density (CPSD) matrix of the full process ***U*** _*t*_. We discuss the latter below, and introduce state-space modelling in the Appendix S2 Appendix: Linear state-space modelling. Efficient MATLAB^©^ implementations of all estimation methods described here are publicly available as open-source software in the Multivariate Granger Causality Toolbox, version 2 (MVGC2 [44, 123]), which may be obtained from https://github.com/lcbarnett/MVGC2.

#### Calculating Granger causality via spectral factorisation

The CPSD of a stationary, zero-mean process ***U***, written *S*(*ω*), is defined as the two-sided Fourier transform of the *autocovariance sequence*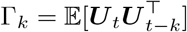, −∞ *< k <* ∞. There are a variety of standard techniques for estimating the CPSD from time-series data, including averaged periodogram, multi-taper, and wavelet methods, or it may be calculated from parametric models.

At any circular frequency *ω* ∈ [−*π, π*], *S*(*ω*) is a Hermitian matrix. Under our assumed regularity conditions, *S*(*ω*) may be uniquely factorised as [124]

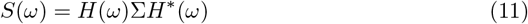

where *H*(*z*), *z* ∈ ℂ is the *transfer function* for the process^13^, superscript ‘*’ denotes matrix conjugate transpose, and Σ is the residual error covariance matrix associated with the optimal least-squares prediction 𝔼 [***U*** _*t*_ | ***U*** _*t*−*τ*_] of ***U*** on its past. Given a CPSD at some frequency resolution, there are stable and efficient algorithms for effecting the *spectral factorisation* (11) [123, 124].

We exploit spectral factorisation to calculate the Granger causality *F* (***X*** → ***Y*** | ***Z***) for a partitioned process 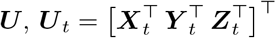 as follows [125]: given the CPSD *S*(*ω*) for the full process ***U***, we obtain the residuals covariance matrix Σ_*yy*_ of (9) by factorising *S*(*ω*) according to (11) and taking the *yy* block of the factored residuals covariance matrix Σ. Next, we note that the *yz* block of *S*(*ω*) is just the CPSD of the restricted process 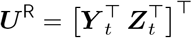; again according to (11), we factorise this restricted CPSD *S*^R^(*ω*) to obtain a restricted residuals covariance matrix Σ^R^, and take the *yy* block to obtain the 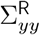 of (9).

We also state here a useful classical result ([126]): under our assumed regularity conditions:

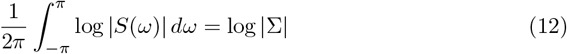

This equation connects the GC estimate obtained through VAR modelling to the spectral factorisation of the CPSD.

#### Dynamical dependence as a Granger causality

We consider the dynamical dependence *T* (***X*** → ***Y***) for an *N* -dimensional microscopic process ***X*** on ℝ^*N*^ and a coarse-grained *n*-dimensional macroscopic variable ***Y*** given by ***Y*** _*t*_ = *M* ***X***_*t*_, where *M* is a full-rank *n × N* matrix. As explained in the main text, without loss of generality we may assume that *M* is an orthogonal matrix: *MM* ^⊤^ = *I*.

We examine the GC for the dynamical dependence, given by

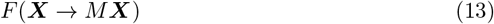

The formula (12) allows us to derive an elegant and computationally-efficient expression for the DD. Firstly, it follows straightforwardly that the restricted CPSD is given by *S*^R^(*ω*) = *MS*(*ω*)*M* ^⊤^, so that we may calculate 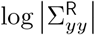 immediately from (12). Secondly, we have Σ_*yy*_ = *M* Σ*M* ^⊤^, so that

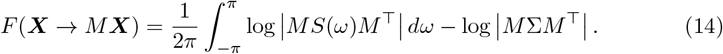

Transformation-invariance of DD allows this to be simplified even further: we can always transform the microscopic space ℝ^*N*^ in such a way as to decorrelate and normalise the residual prediction errors so that Σ = *I*, the identity matrix [30]. Then by orthogonality of *M*, the second term in (14) vanishes, and we obtain

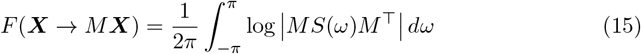

Note that as to the utility of this formula, it does not matter *how S*(*ω*) and Σ have been acquired, e.g., whether it be via parametric modelling or spectral factorisation. It is particularly useful in a DD optimisation scenario: acquisition of *S*(*ω*) and Σ, followed by residuals decorrelation/normalisation, need only be performed once, and (15) may subsequently be applied as often as required for any set of *M*. The computational efficiency of (15) hinges on the dimensionality of *N* and *n*, and in particular the spectral resolution of *S*(*ω*), i.e., the size of the frequency increment Δ*ω* in the numerical quadrature for the integral in (15); the largest acceptable frequency increment will be determined by the complexity of the spectrum of ***X***.

### S2 Appendix Linear state-space modelling

As described in Appendix S1 Appendix: Granger causality, dynamical dependence as a Granger causality may be estimated nonparametrically by estimation of the CPSD and spectral factorisation, or parametrically by vector autoregressive or linear state-space modelling. In this study, we use the linear state-space approach, along with the efficient spectral formula (15) to calculate and minimise DD.

State-space models have a specific advantage over VAR models, with implications for Granger causality estimation: namely, that they are able to parsimoniously model^14^ a moving average (MA) component in the data (they are in fact equivalent to the class of vector-autoregressive moving-average (VARMA) models [107]). This is of particular significance for modelling of neurophysiological data, since common technological and pre-processing factors such as observation noise, downsampling and digital filtering all induce an MA component in the data. Perhaps most importantly, even if a time series is generated by a pure finite-order VAR model, a *sub*-process of that process will generally have an MA component (and any recorded neural process is, of course, inevitably a sub-process of full brain dynamics). The sub-process issue has a particular impact on VAR GC estimation; even if the *full* process is pure, finite-order VAR, the *restricted* process will in general be VARMA. While this is somewhat mitigated by the single-regression VAR estimated mentioned previously, state-space modelling provides a more principled approach, yielding GC estimators of smaller bias and variance. See [44, 127–129] for further discussions on these issues.

We consider observational data to be generated by a discrete-time, continuous-valued, wide-sense stationary stochastic vector process, ***X*** = [*X*^1^, …, *X*^*N*^]^⊤^ on ℝ^*N*^. The linear state-space (SS) model framework assumes that the ***X***_*t*_ are noisy observations of a latent (unobserved) state process ***ξ*** defined on ℝ^*r*^, where the latent space dimension *r* may be larger or smaller than *N*. Specifically,

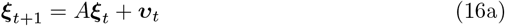

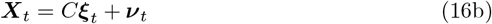

Here *A* is the stable^15^ *r × r* state transition transition matrix, *C* the *N × r* observation matrix, while ***υ*** is a white endogenous noise process, and **ν** a white exogenous (observation) noise process; ***υ*** and **ν** may be contemporaneously correlated, with joint covariance matrix

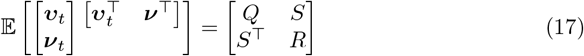

The parameters of the model (16) are *{A, C, Q, R, S}* (the covariance matrix 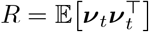 must be positive-definite).

It may be shown that a state-space model in the form (16) may always be transformed to a convenient equivalent model in *innovations form* by introducing the new state variable ***ζ***_*t*_ =𝔼 [***ξ***_*t*_ | ***X***_*t*−*τ*_]. The resulting ISS model takes the form

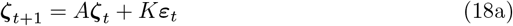

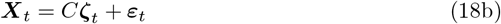

where *K* is the *r × N Kalman gain* matrix, and ***ε***_*t*_ = 𝔼 [***X***_*t*_ | ***X***_*t*−*τ*_] the *innovations* (white noise) process, with covariance matrix Σ = 𝔼 ***ε***_*t*_***ε***^⊤^. The parameters of the model (18) are then *{A, C, K*, Σ*}*, and our regularity conditions (specifically a *minimum-phase* requirement), require stability of *A* − *KC*. We may always transform a general SS model (16) to innovations form (18). We have

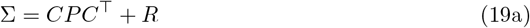

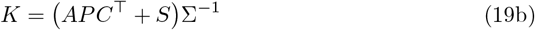

where the *r × r* covariance matrix *P* is the unique stabilising solution of the Discrete-time Algebraic Riccati Equation (DARE)^16^:

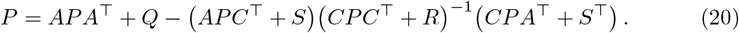

There are efficient and stable computational algorithms available in major programming languages for solution of DAREs, and efficient algorithms too for estimation from time-series data of state-space models in innovations form^17^ [refs].

The transfer function for the ISS model (18) is given by [44]:

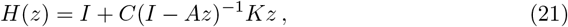

from which the CPSD may be calculated according to (11).

#### Granger causality and dynamical dependence for state-space models

Since the innovations ***ε***_*t*_ in an Innovations-form State Space (ISS) model are just the residual least-squares prediction errors 𝔼 [***X***_*t*_ | ***X***_*t*−*τ*_], the ISS form is well-suited to calculation of Granger causalities [44, 129]. Specifically, if we have an ISS model for a partitioned variable ***U*** = ***X***^⊤^***Y*** ^⊤^***Z***^⊤ ⊤^, then by restricting the observation equation (18b), the restricted process ***U*** ^R^ = ***Y*** ^⊤^***Z***^⊤ ⊤^ is easily seen to follow an SS model, albeit no longer in SS form. We may then solve the appropriate DARE (20) to bring the restricted SS model into innovations form, and in particular to obtain Σ^R^ from (19b). We work through this procedure for dynamical dependence.

Given a state-space model in innovations form (18) and an orthogonal *n × N* linear coarse-graining matrix *M*, from (18b) the observation equation for ***Y*** _*t*_ = *M* ***X***_*t*_ is [44]:

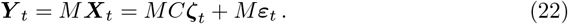

This equation, along with the state-transition equation (18a), constitutes a restricted state-space model, though no longer in innovations form, for which *Q* = *K*Σ*K*^⊤^, *S* = *K*Σ*M* ^⊤^, *R* = *M* Σ*M* ^⊤^ and *C* is replaced by *MC*. We may then solve the corresponding DARE (20) for *P* = *P* (*M*) to convert the restricted SS model to innovations form, which yields, in particular, the restricted residual error covariance matrix

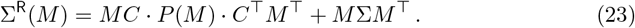

If the full residuals covariance matrix has been decorrelated/normalised to *I*, then this becomes simply

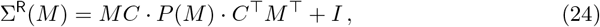

and the dynamical dependence is simply

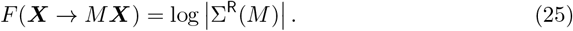

Alternatively, we may calculate the CPSD *S*(*ω*) using the expression (21) for the ISS transfer function, and *F* (***X*** → *M* ***X***) may then be calculated according to the spectral integral (15). Anecdotally, we find that for neurophysiological time-series data, the latter approach is often more computationally efficient than the DARE route. It also facilitates a gradient descent approach to minimisation of dynamical dependence (S3 Appendix).

### S3 Appendix Minimisation of dynamical dependence by gradient descent

A drawback with the DARE method (Appendix S2 Appendix: Linear state-space modelling) is that it is not clear how we might compute the gradient of the DD (25) with respect to the Stiefel parametrisation by orthogonal matrices *M*, since the covariance matrix *P* (*M*) in the expression (24) for Σ^R^(*M*) is defined only implicitly through solution of the restricted DARE. Gradient calculation, though, is indeed possible for the spectral form (15) (Appendix S1 Appendix: Granger causality), regardless of whether the CPSD *S*(*ω*) is acquired through parametric modelling (VAR or SS), or nonparametrically.

We note firstly that since the geometry of the Grassmannian manifold is non-Euclidean, calculation of the DD gradient under the Stiefel parametrisation by orthogonal matrices is not straightforward [67]. For macroscopic scale *n >* 1, furthermore, as remarked previously [ref] the Stiefel parametrisation is not one-to-one, with the implication that there will always be a zero-gradient submanifold in the DD cost surface over the Stiefel manifold^18^. For details of the Grassmannian gradient operator under the Stiefel parametrisation, see [67]; for calculation of the gradient of the spectral-form DD (15), see [30] (APPENDIX D)^19^

A useful approach to speeding up the efficiency of state-space DD optimisation is pre-optimisation of the “proxy” DD:

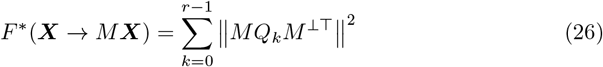

where *M* ^⊥^ is a basis for the orthogonal complement of the hyperplane specified by *M* ^20^, *Q*_*k*_ = *CA*^*k*^*K*, and ∥ *·* ∥ denotes the Frobenius matrix norm. Note that the *Q*_*k*_ depend only on the state-space model parameters, and may thus be precomputed. The proxy DD vanishes at exactly the same points on the Grassmannian as the actual state-space DD, and, as a polynomial in *M*, is computationally much cheaper to evaluate than both the DARE-derived and spectral-form DD. Its gradient may also be calculated explicitly.

Empirically, the local minima of the proxy DD (26) appear in general to be found nearby on the Grassmannian to those of the actual DD. In [30] it is found that preliminary gradient descent using the proxy (26), followed by gradient descent of the spectral-form DD (15), initialised at the optimal hyperplane found by the proxy minimisation run, greatly improves optimisation efficiency.

A simple gradient descent algorithm with adaptive step size is deployed: an initial hyperplane is chosen uniformly at random on the Grassmannian, and the step size set to a specified initial value. At each iteration, a step is taken in the (downhill) direction of the gradient at the current hyperplane, using the current step size, and the DD (or proxy DD) evaluated there. If this DD is smaller than or equal to the current optimal DD, then the new hyperplane becomes the current one (the step is accepted), and the step size is increased by a fixed acceleration factor. Otherwise, the current hyperplane remains unchanged (the step is rejected), and the step size is decreased by a fixed deceleration factor. Optimisation terminates when the step size falls below a specified tolerance, or a maximum number of iterations is exceeded (optimisation times out). Parameters for the algorithm are the initial step size, the acceleration and deceleration factors, the step size termination tolerance, and the maximum number of iterations. See [30] for full details and discussion of alternative optimisation techniques. An open-source MATLAB^©^ implementation of state-space dynamical dependence calculation and minimisation techniques as described here may be obtained from https://github.com/lcbarnett/ssdi.

## S4 Appendix Global brain network dynamics

Globally, the brain network dynamics are governed by an *evolution equation* which takes in the activity of the entire system at time *t* and outputs the vector-valued state of the biophysical neural model at time *t* + 1. Consider a single node *x*_*i*_ within this framework. The dynamical process at node *x*_*i*_ over time is described by the following equation:

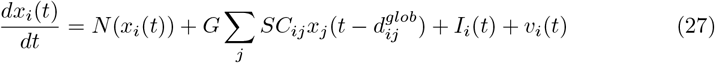

Eq 27 is the same evolution equation used for whole-brain network simulations [9]. Table 1 provides definitions for the terms in Eq 27 as such:

**Table 1.** Evolution equation parameters.

| Parameter | Description |
| --- | --- |
| $\frac{dx_i(t)}{dt}$ | The differential operator mapping system of equations into its derivative i.e. its next time step. |
| $N$ | State operator for each neural mass model, determining the internal dynamics of node. |
| $x_i(t)$ | Vector of state variables for each region (e.g., mean field, or firing rate). |
| $G$ | Global connectivity scaling factor, modulating the influence of global connections on the node's activity. |
| $SC_{ij}$ | Structural connectivity matrix (global coupling in the WBM, at level $\nu = 2$ |
| $d_{ij}^{glob}$ | Time delay between globally coupled nodes, representing the propagation time of signals across the brain network. |
| $I_i(t)$ | External input to the node, representing external stimuli or perturbations affecting the system. |
| $v_i(t)$ | Noise term, accounting for stochastic fluctuations in the system, capturing the inherent variability in neural activity. |

**Table 2.**
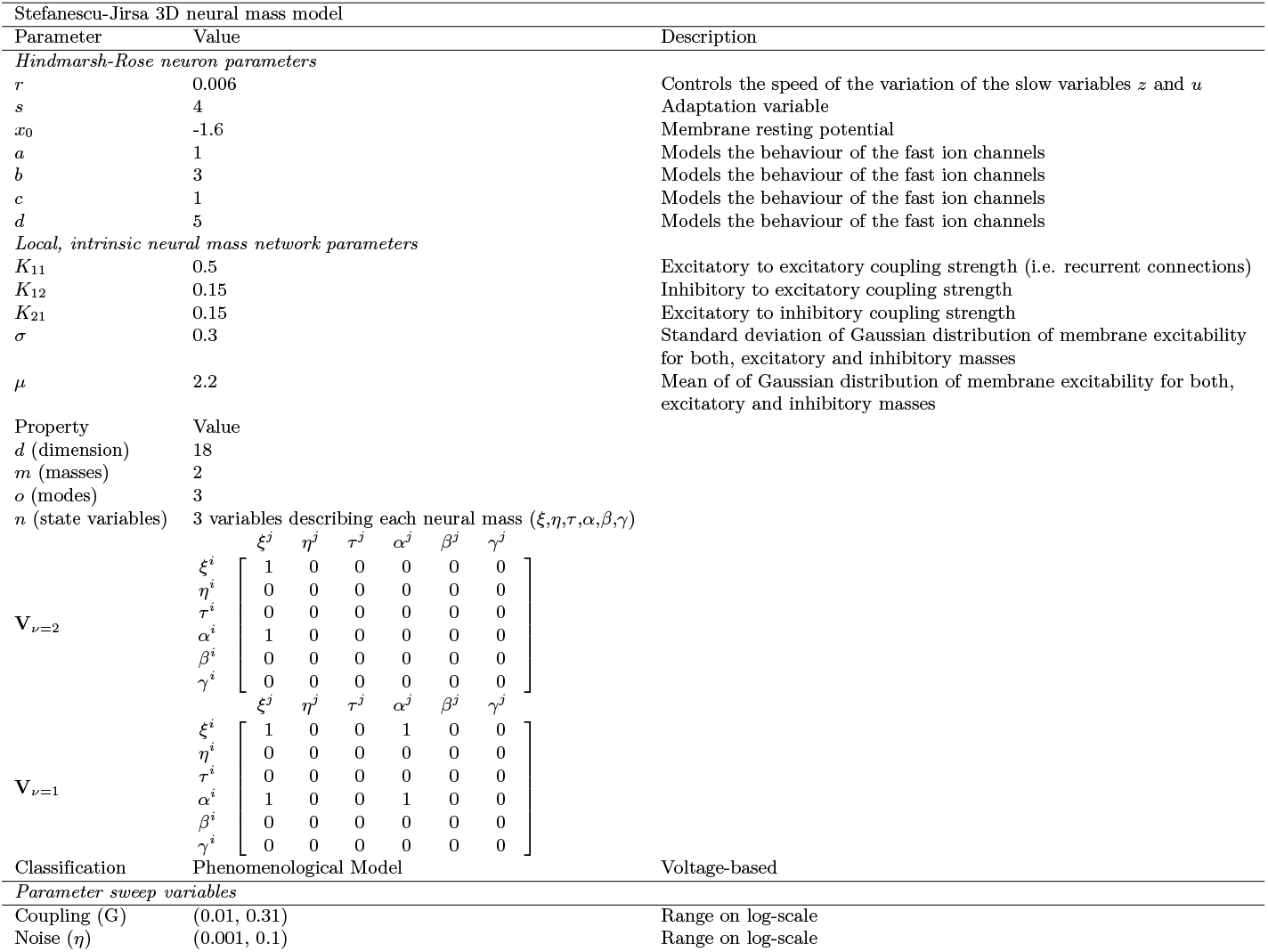
Local Stefanescu-Jirsa 3D and Global Network Parameters.

Two crucial elements in this model are essential. First, temporal delays are included on global connections, providing a more realistic understanding of how brain activity propagates at specific nodes [7, 72, 130]. These delays are particularly critical at the WBM level, where interactions between localised brain areas unfold over time on the mesoscopic level, leading to the large-scale, macroscopic, spatiotemporal dynamics observed in the whole brain. Second, a veridical structural connectivity matrix implements the large-scale network. This matrix is usually obtained from magnetic resonance imaging (MRI) scans.

In summary, WBMs are versatile, accommodating both local interactions within cortical areas and global interactions across the entire brain. This allows researchers to model how neural dynamics evolve both locally and globally, providing insights into the mesoscopic processes that give rise to patterns of macroscopic activity.

### Neural mass model

The Stefanescu-Jirsa 3D (SJ3D) model is derived from a mean-field approximation of coupled Hindmarsh-Rose (HR) neuronal models. HR neurons are excitable neuronal models that are capable of exhibiting neuronal bursting oscillations, endowing the subsequent SJ3D model with capability of displaying qualitatively complex dynamics such as synchronisation, metastability, multi-clustering, spike-burst behaviour, oscillatory death and spiking limit-cycle oscillations [9, 52, 53].

The HR model consists of three coupled differential equations:

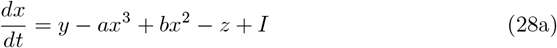

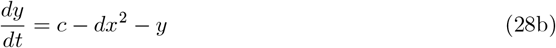

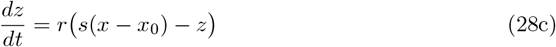

Here, *x* and *y* are state-variables that evolve on a faster time-scale, dictating the spike-burst activity, and *z* is a state-variable representing the slower time-scale changes usually representative of the oscillatory activity. *I* is an external input parameter and is responsibly for altering the system dynamics from fixed-point dynamics (*I <* 1.3), spike-bursting behaviour (*I >* 1.3). Which additionally, can be expressed in a chaotic regime or an oscillatory regime.

Given that we want to describe two neural masses (populations) within one neural mass model, we can therefore adapt the HR model for population dynamics as such:

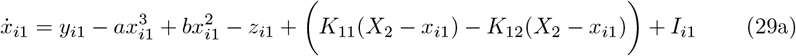

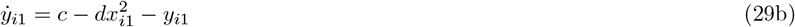

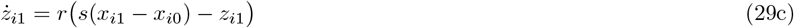

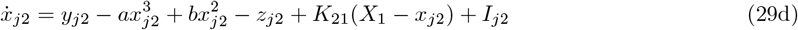

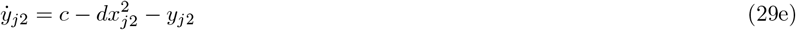

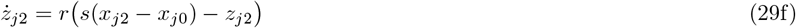

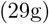

Above, the mesoscopic coupling between excitatory and inhibitory neural masses is indicated by the *K*_*nm*_ parameters and is given in Table 2. Given mode decomposition methods as presented in Assisi *et al*. [70], the following reduced equations are derived by the original authors [53]:

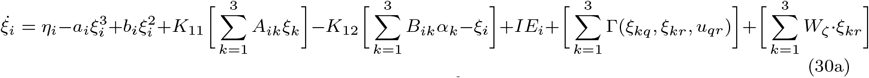

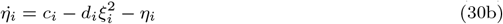

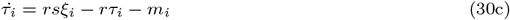

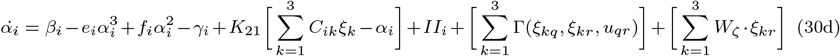

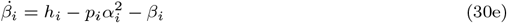

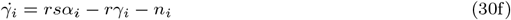

Now, the connectivity on the macroscopic scale of brain regions is given introduced by the *W*_*ζ*_ parameter. The full description of the parameters adapted from [69] is given in Table 2. For the interested reader, the comparison between the full model (Eq 29 and the reduced equations (Eq 30) is performed in the original articles [52, 53].

Table 2 also includes the range of global coupling and dynamical noise for the bivariate parameter sweep. Moreover, **V**_ν=2_. Here, following the original notation of Sanz-Leon [9], **V**_ν=2_ corresponds to the inter-regional connectivity between brain regions or nodes, and **V**_ν=1_ corresponds to the intra-regional connectivity between excitatory and inhibitory populations with the individual NMMs.

## S5 Appendix Principal angles & single-node contribution to macroscopic dynamics

Principal angles between subspaces, specifically macroscopic variables of dimension *n*, can be used to define invariant metrics on the corresponding Grassmannian manifold [131]. Formally, given two subspaces in ℝ^*n*^ a metric on the Grassmannian can be defined as 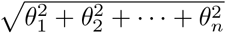, where each *θ*_*i*_ ∈ (0, *π/*2) represents a principal angle corresponding to the coordinate axes of an *n*-dimensional macroscopic variable.

Hereafter, these macroscopic variables are denoted as *n*-macros.

To quantify the similarity between two *n*-macros of the same dimension, obtained from different optimisation runs with random initial restarts, we define a metric that is normalised within the interval [0, 1]. Here, 0 represents complete coplanarity (generalised colinearity), while 1 indicates orthogonality between the *n*-macros. Specifically, for a *n*-macro, the similarity metric is given by 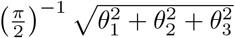, where perfect coplanarity (identical *n*-macros) results in a metric value of 0.

This metric forms the foundation for identifying the *emergent dynamical structure* associated with higher-order interactions across spatial scales. By exploiting the optimisation landscape’s structure—specifically, the geometry of the Stiefel manifold—we determine whether local minima dominate the energy landscape. Since these local minima correspond to *n*-macros, they provide a proxy for assessing the dynamical dominance, and hence, the emergent dynamical structure at higher-order scales.

Similarly, using principal angles, we can evaluate the extent to which the dynamics are *localised* within the original microscopic state-space.

### Localisation of *n*-macros to microlevel nodes: a single-node approach

Given the identified emergent *n*-macros and their corresponding DD values, a natural question arises: *Which microlevel nodes contribute to the dynamics of the n-macro, and to what extent* ? This consideration enables the projection of *n*-macros back onto the microscopic base, or causal graph, to determine where the macroscopic dynamics are localised within the original microscopic state space. This provides a specific perspective on the *emergent dynamical structure* by the degree of distinct localisation of the macroscopic processes on the constituents of the microscopic process.

Similarly to measuring the principal angles between subspaces (*n*-macros) on the same spatial scale, we can measure the principal angles between the coordinate axes of the *N* -dimensional microscopic base and a candidate *n*-macro. This generalisation of principal angles applies to any two subspaces of arbitrary dimensions. The degree of single-node contribution is now represented by 1− the metric distance: 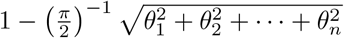. In this case, a value closer to 1 indicates a higher degree of single node contribution to the *n*-macro.

To formally define what it means for a microscopic node to contribute to the *n*-macro, consider a system of three variables *{X*^1^, *X*^2^, *X*^3^*}* undergoing a linear transformation induced by an emergent 2-macro at time *t*. This can be expressed as the matrix multiplication:

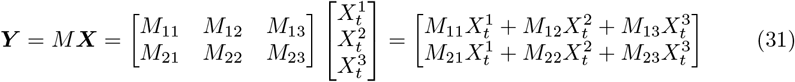

Here, *N* and *n* represent the dimensions of the microscopic base and the *n*-macro, respectively. Each element *M*_*nN*_ parameterises the 2-macro and corresponds to the element in the Stiefel manifold represented by the orthonormal matrix *M*, determining the contribution of each microscopic node 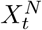 to the macroscopic process. For instance, if *M*_11_ = *M*_21_ = 0, the macroscopic dynamics governed by the 2-macro are entirely driven by *X*^2^ and *X*^3^. In the context of a brain network, where *X*^2^ and *X*^3^ represent two regions in a three-region network, this indicates that these two regions contribute to the emergent 2-macro, while the *X*^1^ region does not.

Collectively, the 2-macro and the single node *X*^1^ encapsulate the the localisation of the macroscopic variable, and therefore a perspective of the emergent dynamical structure of this simplified system. This provides a dimensionally reduced description of the dynamics without resorting to the full higher-dimensional microscopic dynamics. The free parameters *M*_*nk*_ define the principal angle between any *n*-macro and the microscopic base ***X***.

### Localisation of *n* -macros to microlevel nodes: a grouped-node approach

In higher-dimensional systems, the relationship between the coordinate axes of *n*-macros and those of the microscopic system becomes more complex, as the specificity of which nodes contribute to the *n*-macro is harder to determine. Formally, this reflects the fact that the set of subspace angles (principal angles) between an *n*-macro and the coordinate axes of the microlevel constituents does not uniquely identify the *n*-macro. In higher dimensions, distinct subspaces (*n*-macros) can share the same set of angles with the coordinate axes, and can sometimes lead to ambiguity in pinpointing the contributing microlevel nodes [30].

Given the high dimensionality of neurophysiological recordings or region-based whole-brain models—often encompassing between 32 and 256 variables—subspace angles between each individual node of the microscopic process and *n*-macros may not uniquely determine the localisation of the emergent dynamical structure in the microscopic state-space. However, despite these challenges, assessing the contribution of individual nodes to *n*-macros remains computationally tractable and highly informative, especially when a researcher seeks a fine-grained understanding of the microlevel localisation of the *n*-macros.

As an alternative, we could impose an additional constraint by measuring the subspace angle between each emergent *n*-macro and all *n*-combinations of microlevel variables. Formally, this is defined as the subspace angle between the emergent *n*-macro and all *n*-subsets of the original *N* -dimensional microlevel state-space, thus there are 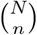 such combinations. Unlike angles with coordinate axes, these 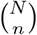 subspace angles do uniquely specify the subspace—though at the cost of a combinatorial explosion. While this approach offers less specificity than measuring the subspace angle between each node and *n*-macro, it serves as a proxy for partitioning the system into distinct emergent cores.

The *n*-dimensional subset that is optimally implicated in driving the macroscopic dynamics is identified as the *n*-subset with the minimal principal angle from the *n*-macro. As previously discussed, though this is illustrated here, this alternative approach is not utilised in empirical data within this thesis particularly because of its partitioning nature.

## S6 Appendix Worked example

This section illustrates the pipeline for implementing DI on stationary discrete-time, continuous-valued multivariate stochastic processes using a linear state-space (SS) parameterisation, details of which can be found in S2 Appendix: Linear state-space modelling. The general computational routine is schematically shown as a geometric interpretation with corresponding data visualisations in Fig 8.

First, we simulate synthetic data from a 9-variable VAR(9) model, where the temporal model order (i.e., the number of discrete-time lags considered in the prediction) is *p* = 8. The model was prespecified with specific structural connectivity (VAR coefficients) that reflect the G-causal structure evident in Fig 9 We then compute the full causal graph representing the directed information flow between pairs of variables (Fig 9). This causal graph serves as the *functional* ground truth model of the microlevel dynamics and the information flow between the variables. The variables in our system correspond to the activity of network nodes in the causal graph. Therefore, without loss of generality, we will refer to these variables as nodes from this point forward.

**Fig 9.**
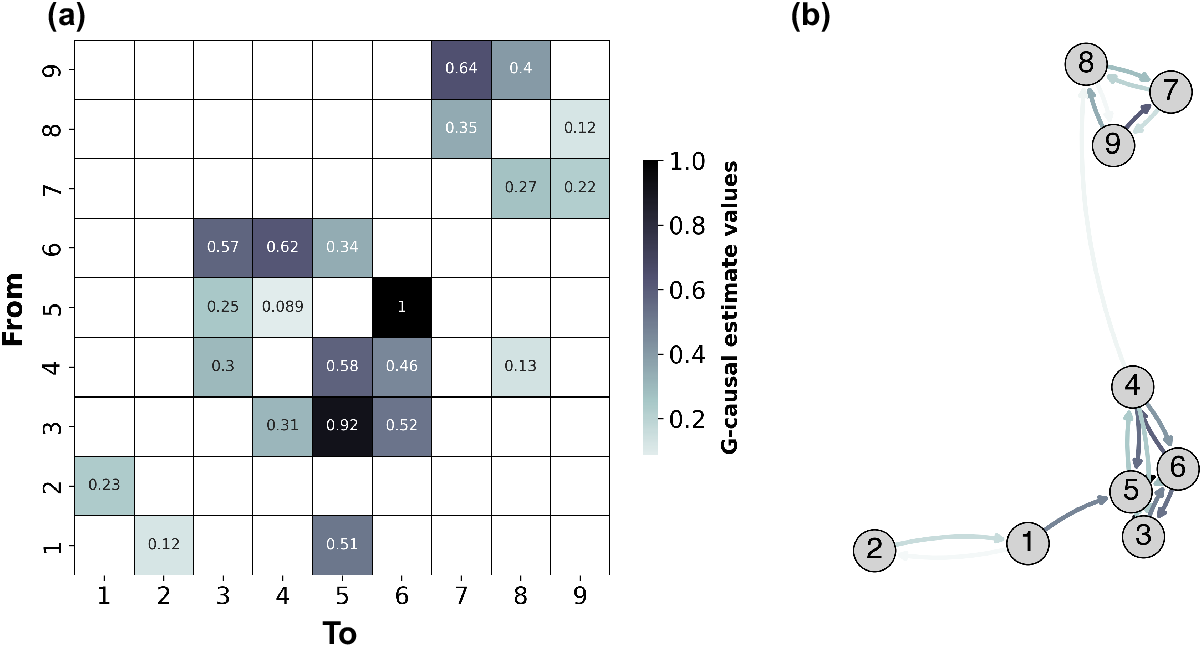
Pairwise Conditional Causal Graph of a VAR(9). The figure illustrates the strength and directionality of connections between variables in a 9-variable VAR model, with darker colors indicating stronger causal relationships. (a) Matrix representation of the pairwise causal estimates, where each cell’s colour intensity reflects the strength of the causal effect from the variable on the y-axis (”From”) to the variable on the x-axis (”To”). (b) Graph representation of the causal relationships, where the arrow colors correspond to the same weighting as in the matrix, visually depicting the directional influences between variables.

All pairwise directed information flow estimates were retained to avoid pruning the causal graph based on statistical significance. Pruning, which is commonly done to retain only significant directed functional connections, would risk disrupting the statistical structure of the entire system. By keeping all pairwise estimates, we preserve the full range of statistical information inherent in the system. This approach assumes that information that may not be apparent at the microscopic scale could still contribute to the identification of the emergent macroscopic dynamical structure.

We employ a pre-optimisation step to accelerate computational efficiency (see S3 Appendix: Minimisation of dynamical dependence by gradient descent for details). This step performs 100 random restarts, which terminate either after 10,000 iteration steps or if the gradient falls below the threshold set at 1*e*^−10^. Then, *n*-macros are clustered according to their pre-optimisation DD values and the optimisation of DD is performed on the reduced clusters. The optimisation history panels in Fig 10 illustrate the remaining optimisation runs after pre-optimisation clustering for a *n*-macros of scale 2 to 8 whilst the similarity matrices in Fig 10 illustrate the similarities between the *n*-macros across optimisation runs.

**Fig 10.**
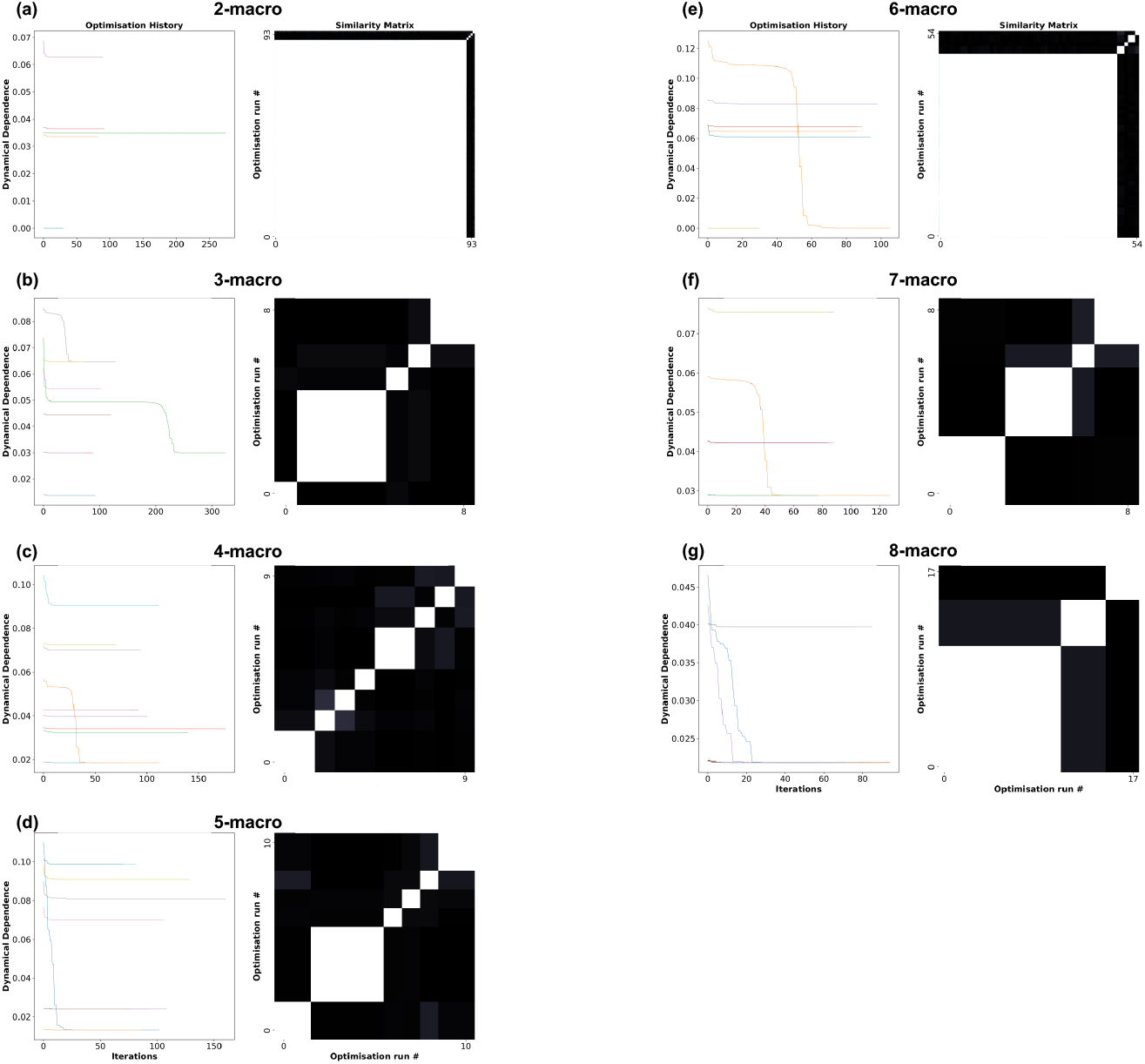
Optimisation Histories and Similarity Matrices for *n* -macros. Panels (a)-(g) present the optimisation histories and corresponding similarity matrices for coarse-grainings (*n*-macros), where *n* ranges from 2 to 8. The optimisation panels display the results *after* clustering, following the initial pre-optimisation procedure. The similarity matrices illustrate the degree of similarity between local minima (i.e., different *n*-macros) identified across optimisation runs with random restarts. White indicates identical *n*-macros, while darker shades represent increasingly dissimilar *n*-macros. We see that only a 2-macro and 6-macro converge onto a value of absolute dynamical independence (value of zero)

Results show that distinct optimisation runs converge to distinct values of DD. Def 3 states that an emergent *n*-macro is the *minimally* dynamically dependent macroscopic variable at the prescribed spatial scale. We observe that given the prespecified underlying structure of the causal graph (Fig 9) an observer might preempt the possibility of discovering perfect dynamical *independence* (Eq 2) for both; a 2-macro, 3-macro, and 4-macro. However, results show that this is only identified for a 2-macro and a 6-macro (Fig 10 panels(a) and (e)).

Given that dynamical dependence converges to different values across the optimisation runs, it raises the question: *Are the n-macros corresponding to these different dynamical dependence values the same?* The similarity matrices in Fig 10(Similarity matrices panels) provide insight into this question by illustrating the similarity between *n*-macros identified across 100 optimisation runs on the same scale, *n*. The similarity between *n*-macros is quantified by the principal angle between them (see Section).

The optimisation results presented in Fig 10 show that runs converging to the same dynamical dependence value correspond to the same *n*-macro, as evidenced by the high similarity in the matrices (White cells in the Similarity matrices panels of Fig 10). In contrast, when optimisation runs converge to different dynamical dependence values, the corresponding *n*-macros exhibit varying degrees of dissimilarity^21^. Furthermore, the results indicate a predominant single 2-macro structure within the optimisation landscape, whereas the 3-macro results display greater diversity, suggesting a more complex landscape with multiple distinct local-minima.

These similarity matrices indicate the presence of basins of attraction in the energy landscape suggesting that a single *n*-macro can dominate the dynamics at a particular scale. This dynamical dominance indicates that the randomised optimisation restarts (the gradient descent) consistently converge to the same local minima (*n*-macro); reflecting a strong emergent dynamical structure at that scale. Such basins of attraction offer a powerful way to assess another aspect of emergence structure—specifically, the stability and uniqueness of the emergent dynamical structure at higher-order scales. Although this method of evaluating emergent dynamics is not utilised in the following chapter, it provides a promising direction for future research, particularly in understanding how these basins of attraction contribute to the formation of stable, higher-order structures.^22^

When dealing with high-dimensional systems, single-node contributions (Section) may always meet the necessary empirical criteria, as subspace angles between emergent *n*-macros and their microlevel constituents do not generally provide a unique identification of the macroscopic subspace. To address this limitation, a grouped-node approach offers a valuable alternative, building on the subset classification illustrated in Fig 11. However, it is important to note that using grouped-node level analysis involves a trade-off between capturing the nuanced, graded nature of localised contributions and inducing a partitioning of the system. Therefore, the choice of analysis should be carefully considered and guided by the specific empirical question being addressed. For details on the full implementation, see Section. Here we offer both approaches for completeness.

**Fig 11.**
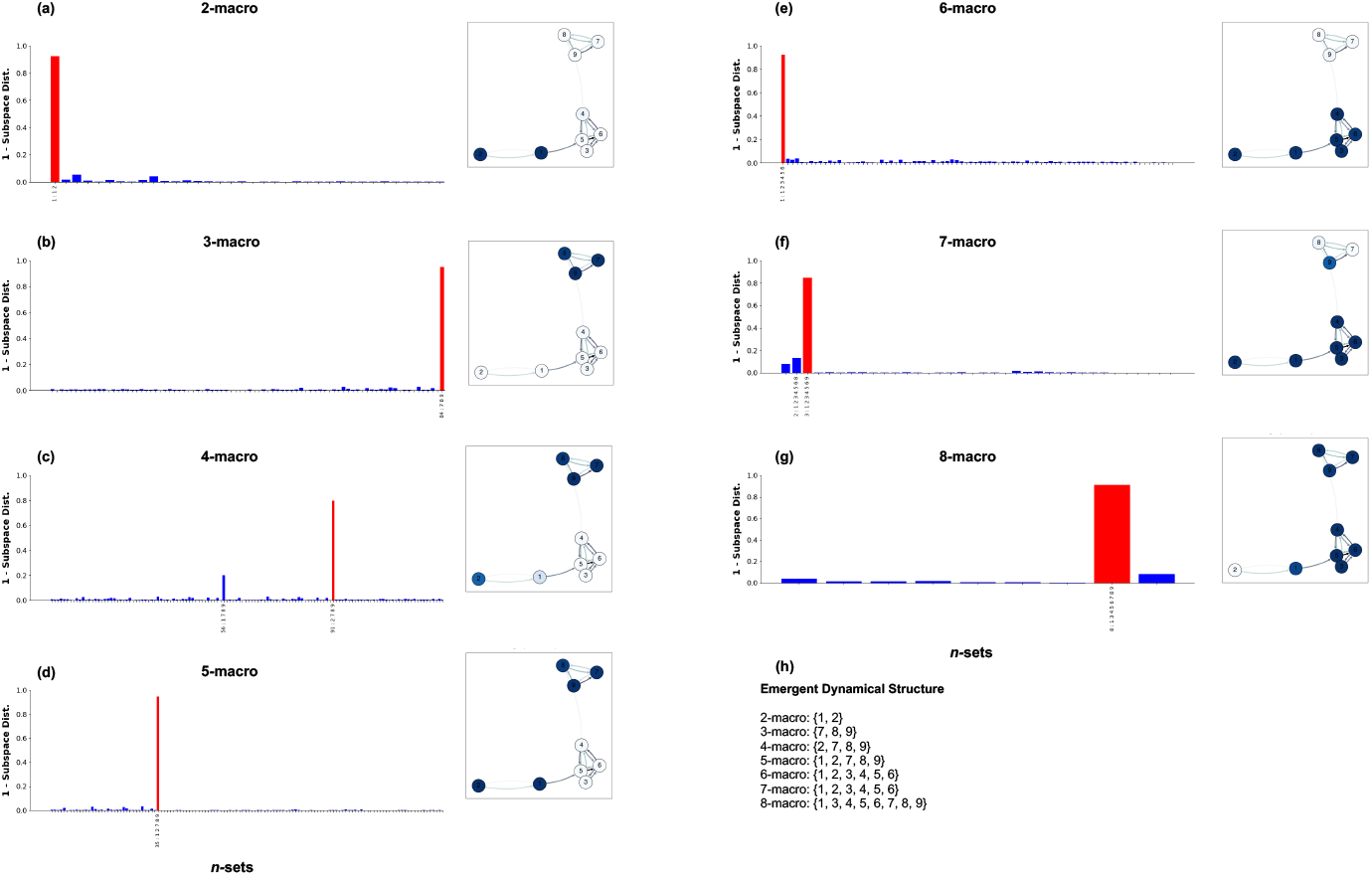
Single-node and Grouped-node Contributions to Emergent *n* -Macros. Panels (a-g) show the contributions of individual nodes and grouped-nodes (see Section) to emergent *n*-macros, ranging from 2-macros to 8-macros: Histograms represent grouped-node contributions, where red bars indicate the *n*-subsets of microlevel nodes that are maximally aligned with the *n*-macro. Only contributions greater than 0.1 are shown. Inset causal graphs depict single-node contributions, with node sizes reflecting their individual contribution on the *n*-macro. Panel (h) summarises the distinctly partitioned subsets using grouped-node analysis that optimally contribute to each *n*-macro across different scales.

Fig 11 explores the contributions of individual and grouped nodes to emergent *n*-macros across a range of spatial scales, from 2-macros to 8-macros. Each panel (a-g) includes a histogram and an inset causal graph with the emergent *n*-macro projected onto it, offering complementary perspectives on how microlevel nodes contribute to the emergent dynamics.

The histograms illustrate grouped-node contributions, where each bar represents the degree of co-planarity between a specific *n*-subset of nodes and the *n*-macro. A red bar highlights the subset that is most strongly aligned with the *n*-macro, indicating the microlevel nodes that are most integral to the emergent structure. Contributions are only displayed for subsets with values exceeding 0.1, emphasising the most significant relationships.

The inset causal graphs provide a visual representation of single-node contributions, mapping these contributions back onto the original causal graph. In these graphs, the colour correspond to the strength of the contribution to the *n*-macro, with darker colors indicating stronger connections.

Finally, panel (h) offers a summary of the emergent dynamical structure across all scales, listing the specific node subsets that optimally contribute to each *n*-macro. In this sense it partitions the system without a graded analysis of contribution offered by the single-node analysis. This analysis reveals the complexity and high-order nature of dynamical structures in the system, where both individual and collective node contributions can be implemented. This concludes our worked example.

To understand the emergent dynamical structure by assessing how *n*-macros are localised in space across nodes, we aim to avoid inducing a partitioning of node contributions to *n*-macros. Therefore, we utilise the more sensitive and nuanced single-node analysis in our biophysical neural model.

## Acknowledgments

We would like to thank the Melbourne School of Psychological Sciences at the University of Melbourne, the Sussex Centre for Consciousness Science at the University of Sussex, and the Paris Brain Institute (ICM) for providing the necessary resources and institutional support. We also extend our gratitude to lab members and colleagues for their insightful discussions and assistance over the drafting of the manuscript.

This research was supported through a National Health and Medical Research Council (NHMRC) grant #APP1183280 and internal University of Melbourne research funding of Prof. Olivia Carter. The research was also supported, in part, by the European Research Council (ERC) Advanced Investigator Grant 10109254 to A.K. Seth, which also supports L. Barnett. We also acknowledge the Australian Government Research Training Program Scholarship, which made this work possible. Additionally, we appreciate the use of The Virtual Brain (TVB) software, which was crucial to the biophysical neural model simulations and analyses in this study.

Finally, we thank Adam Barrett, Marcello Massimini, Fernando Rosas, Pedro Mediano, and Aniko Kusztor for invaluable discussions, encouragement, and ongoing support throughout the research process.

## Author Contributions

- **Conceptualisation**: Borjan Milinkovic, Thomas Andrillon, Lionel Barnett, Anil K. Seth, Olivia Carter.
- **Formal analysis**: Borjan Milinkovic.
- **Funding acquisition**: Olivia Carter, Anil K. Seth.
- **Investigation**: Borjan Milinkovic.
- **Methodology**: Borjan Milinkovic, Lionel Barnett.
- **Supervision**: Olivia Carter, Thomas Andrillon, Lionel Barnett, Anil K. Seth.
- **Writing – original draft**: Borjan Milinkovic.
- **Writing – review & editing**: Borjan Milinkovic, Thomas Andrillon, Lionel Barnett, Anil K. Seth, Olivia Carter.

## Code Availability

The code used in this study integrates multiple toolboxes and custom scripts. The core method for Dynamical Independence (DI) analysis is implemented through the **SSDI toolbox** developed by Lionel Barnett, which is available at: **https://github.com/lcbarnett/ssdi**.

This toolbox depends on the **MVGC2 toolbox**, also developed by Lionel Barnett, for multivariate Granger causality analysis, available at: **https://github.com/lcbarnett/MVGC2**.

Additionally, the simulations of biophysical neural models were adapted from **The Virtual Brain (TVB)** software, which is available at: **https://www.thevirtualbrain.org/**.

A custom package specifically developed for this project is available on the first author’s GitHub under the repository **TVBEmergence**: **https://github.com/bmilinkovic/TVBEmergence/**.

This package includes all the scripts necessary to reproduce the results, including instructions for environment setup and dependencies.

For further inquiries or assistance with the code, please contact the corresponding author.

## Footnotes

1 The linear mapping is surjective iff *M* has full rank *n*.

2 we use the notation *t* − *τ*, where *τ* is not a single value but represents a set of lagged time points, *τ* = 1, 2, 3, …, *n*, referring to the history of the process. In the case of an infinite history *τ* = 1, 2, 3, …, this set would extend indefinitely.

3 We note, for example, that dynamical independence is *transitive* [30], and that emergent macrovariables may thus potentially (but not necessarily) be nested, leading to the possibility of heterarchical emergence structures.

4 The subspace associated with a coarse-graining is spanned by the *n* basis vectors defined by the columns of the *N × n* transposed coarse-graining matrix *M* ^⊤^. Exploiting the invariance of DD under transformations of the macroscopic space ℝ^*n*^, it turns out to be convenient to parametrise the Grassmannian by *orthogonal* matrices *M* ; i.e., matrices for which *MM* ^⊤^ = *I*. These matrices constitute the *Stiefel manifold* of orthonormal bases, written 𝒱 _*N*_ (*n*)—also a homogeneous Riemannian manifold— and we consequently refer to the parametrisation of the Grassmannian 𝒢 _*N*_ (*n*) by orthogonal *n × N* matrices as the Stiefel parametrisation. We note that for *n >* 1 this parametrisation is not one-to-one; 𝒱 _*N*_ (*n*) has dimension 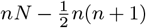, which for *n >* 1 is larger than the dimension *n*(*N* − *n*) of 𝒢 _*N*_ (*n*). In practice, optimising over the Grassmannian and Stiefel manifolds produce similar results [66], and the latter is simpler to implement.

5 Computational neuroscience has a tendency to present functional structure in terms of networks. We stress that in DI, macroscopic variables are, in general, *not* networks in any meaningful sense.

6 In higher-dimensional systems, the axis angles do not uniquely identify a single subspace: the higher dimensionality affords more degrees of freedom for subspaces. Nonetheless, axis angles are informative about region contribution.

7 the analysis for the emergent 2-macros shows exactly similar varied pattern, excluded here for brevity

8 More precisely, any finite subset of process variables at arbitrary time stamps has a multivariate-normal distribution.

9 The generalised variance of a random vector is the determinant of its covariance matrix.

10 Besides stationarity, the GC formalism requires some further regularity conditions on the process ***U*** ; for all stochastic processes in this study we assume the same conditions as described in [119, §2, p. 305].

11 On the analytical level, prediction is based on the *infinite* past. For empirical data, histories will of necessity be truncated at some suitable number of lags.

12 The null hypothesis of vanishing GC may also be tested by an *F* -test, which anecdotally is more powerful than the 𝒳 ^2^ test, especially for small samples

13 By abuse of notation, we write *H*(*ω*) for *H*(*z*) with *z* = *e*^−*iω*^ on the unit circle in the complex plane.

14 That is, with lower model complexity.

15 Our regularity conditions imply stability of *A*, which means that all its eigenvalues lie strictly within the unit circle in the complex plane.

16 The DARE arises as the steady-state limit of the predictive Kalman filter for the SS model (16) [129].

17 We recommend in particular the various “state-space-subspace” algorithms, which are non-iterative and yield quasi-log-likelihood estimates for the model parameters.

18 While there are indeed one-to-one parametrisations of the Grassmannian, they are inhomogeneous and not well-suited to implementation of gradient descent. In practice, however, the redundancy inherent in the Stiefel parametrisation does not appear to be a major impediment for gradient descent.

19 We remark that it is also possible to calculate the appropriate *Hessian* (2nd-order differential operator) under the Stiefel parametrisation of the Grassmannian, and hence for the spectral-form DD, which in principle opens the way to 2nd-order gradient techniques such as Newton’s method or conjugate gradient methods [67]. We found, though, that the computational costs of the Hessian calculation outweighed any gain in optimisation efficiency. We thus confine ourselves here to 1st-order methods.

20 This may be calculated via a Singular Value Decomposition (SVD) of *M*.

21 though in the figure here the dissimilarity seems to be general substantial between macros given the darkness of the cells, in general, this will not be the case, and dissimilarity can be significantly graded

22 This approach is applied and explored in more detail in Chapter 5 to assess the emergent dynamical structure across higher-order scales associated with global states of consciousness

